# EVI/WLS function is regulated by ubiquitination and linked to ER-associated degradation by ERLIN2

**DOI:** 10.1101/2020.12.09.417667

**Authors:** Lucie Wolf, Annika Lambert, Julie Haenlin, Michael Boutros

## Abstract

Wnt signalling is important for development in all metazoan animals and associated with various human diseases. The Ubiquitin-Proteasome System (UPS) and regulatory ER-associated degradation (ERAD) has been implicated in the secretion of WNT proteins. Here, we investigated how the WNT secretory factor EVI/WLS is ubiquitinated, recognised by ERAD components, and subsequently removed from the secretory pathway. We performed a focused, immunoblot-based RNAi screen for factors that influence EVI/WLS protein stability. We identified the VCP-binding proteins FAF2 and UBXN4 as novel interaction partners of EVI/WLS and showed that ERLIN2 links EVI/WLS to the ubiquitination machinery. Interestingly, we found in addition that EVI/WLS is ubiquitinated and degraded in cells irrespective of their level of WNT production. K11, K48, and K63-linked ubiquitination is mediated by the E2 ubiquitin conjugating enzymes UBE2J2, UBE2N, and UBE2K, but independent of the E3 ligases HRD1/SYVN. Taken together, our study identified factors that link UPS to the WNT secretory pathway and provides mechanistic details on the fate of an endogenous substrate of regulatory ERAD in mammalian cells.

## INTRODUCTION

Cell-cell communication is fundamental to multicellular organisms and as such requires tight and nuanced regulation on many levels. Protein stability and turnover are essential to guarantee flexible and context-dependent cellular responses to signalling cues, beside various other mechanisms. Eukaryotic protein degradation is mediated by two main systems: the ubiquitin-proteasome system (UPS) and the autophagy-lysosomal pathway, which can both be initiated by tagging substrates with the small protein ubiquitin (Pohl & Dikic, 2019). The specificity of this post-translational modification relies on the subsequent action of three different enzymatic processes, namely the activation of ubiquitin by the enzyme E1, followed by its transfer to an ubiquitin conjugating enzyme E2 and to the target polypeptide with the help of an ubiquitin ligase E3 (Swatek & Komander, 2016). Whereas substrate recognition is the main task of the numerous E3 ubiquitin ligases, only approximately 35 mammalian E2 conjugating enzymes regulate the initial priming with single ubiquitin moieties and the formation of polyubiquitin chains (Deol et al., 2019; van Wijk et al., 2009; Weber et al., 2016). Since ubiquitin itself has eight potential acceptor sites for further ubiquitin modifications (Lysine (K) 6, K11, K27, K29, K33, K48, K63, and its N-terminus Methionine 1), the resulting chains can be of variable geometry and length (Clague et al., 2015; Deol et al., 2019). K48- and K63-ubiquitin linkage are the best studied and most common types and are functionally primarily involved in proteasomal degradation (K48) or proteasome independent processes (K63), e.g. endosomal trafficking and targeting to the lysosomes, respectively (Akutsu et al., 2016; Clague et al., 2015; Erpapazoglou et al., 2014; Swatek & Komander, 2016).

A major role for the UPS is to remove terminally misfolded proteins to prevent their aggregation and potential harm to the cell, even from within the endoplasmic reticulum (ER). ER-associated degradation (ERAD) recognises, extracts, ubiquitinates, and delivers ER membrane proteins or proteins within the secretory route to the proteasome (Bhattacharya & Qi, 2019; Christianson & Ye, 2014; Lopata et al., 2020; Sun & Brodsky, 2019). Several ER membrane resident E3 ubiquitin ligases provide the poly-ubiquitin signal for ERAD, most notably HRD1, GP78, and MARCH6, which have well-studied orthologs in yeast (Lopata et al., 2020; Preston & Brodsky, 2017). Another major protein in this process is the AAA ATPase VCP/p97 (Cdc48 in yeast), which uses ATP to extract substrates from the ER or its membrane into the cytoplasm and can be recruited to the ER membrane by proteins with VCP binding domain, such as FAF2 and UBXN4 (Bodnar & Rapoport, 2017; Meyer & Weihl, 2014).

Beside the regulation of protein quality control, ERAD can also impact on cellular signalling by regulating the availability of mature proteins through quantity control (Bhattacharya & Qi, 2019; Hegde & Ploegh, 2010; Printsev et al., 2017). It is assumed that ERAD quality and quantity control are mechanistically similar and differ mostly during substrate recognition (Hegde & Ploegh, 2010). While misfolded proteins can be recognised by exposed hydrophobic patches or prolonged retention within the ER, the selection of properly folded and functional proteins for degradation is less straightforward and was only investigated in detail for very few mammalian substrates. Indeed, only about 20 to 30 endogenous substrates for mammalian regulatory ERAD have been reported and for most it is unclear how they are recognised, ubiquitinated, and linked to the ERAD machinery (Bhattacharya & Qi, 2019; Printsev et al., 2017). Furthermore, various studies focus on ubiquitination by HRD1 while other ER membrane resident E3 ligases remain poorly defined (Fenech et al., 2020).

Protein stability is at the heart of many cellular communication pathways and it is not surprising that signalling cascades have numerous intersections with the UPS. An example is the regulated degradation of β-catenin in the absence of active WNT signalling (Aberle et al., 1997), a pathway that controls embryonic growth and patterning and ensures tissue homeostasis in adults, while its deregulation can lead to cancer (Nusse & Clevers, 2017; Zhan et al., 2017). In line with these findings, we recently demonstrated that the conserved transmembrane protein EVI/WLS is a target of regulatory ERAD and is ubiquitinated by the E2 enzyme UBE2J2 and the E3 ligase CGRRF1, before being removed from the ER with the help of VCP (Glaeser et al., 2018). EVI/WLS degradation is inhibited by its binding to WNT ligands, which are modified with palmitoleate by the acyl-transferase Porcupine (PORCN) (Takada et al., 2006). However, it remains unclear how EVI/WLS is linked to the ubiquitination machinery and how VCP is recruited, as EVI/WLS itself has no VCP interaction domain (Figure 1A). Furthermore, knock-down of UBE2J2 and/or CGRRF1 did not completely abolish EVI/WLS ubiquitination, indicating the involvement of additional E2 and/or E3 enzymes, which are currently unknown (Glaeser et al., 2018). In this context, ERAD of EVI/WLS seems to be especially interesting due to its independence of the well-studied ERAD associated E3 ligases HRD1, GP78, and MARCH6.

**Figure 1.**
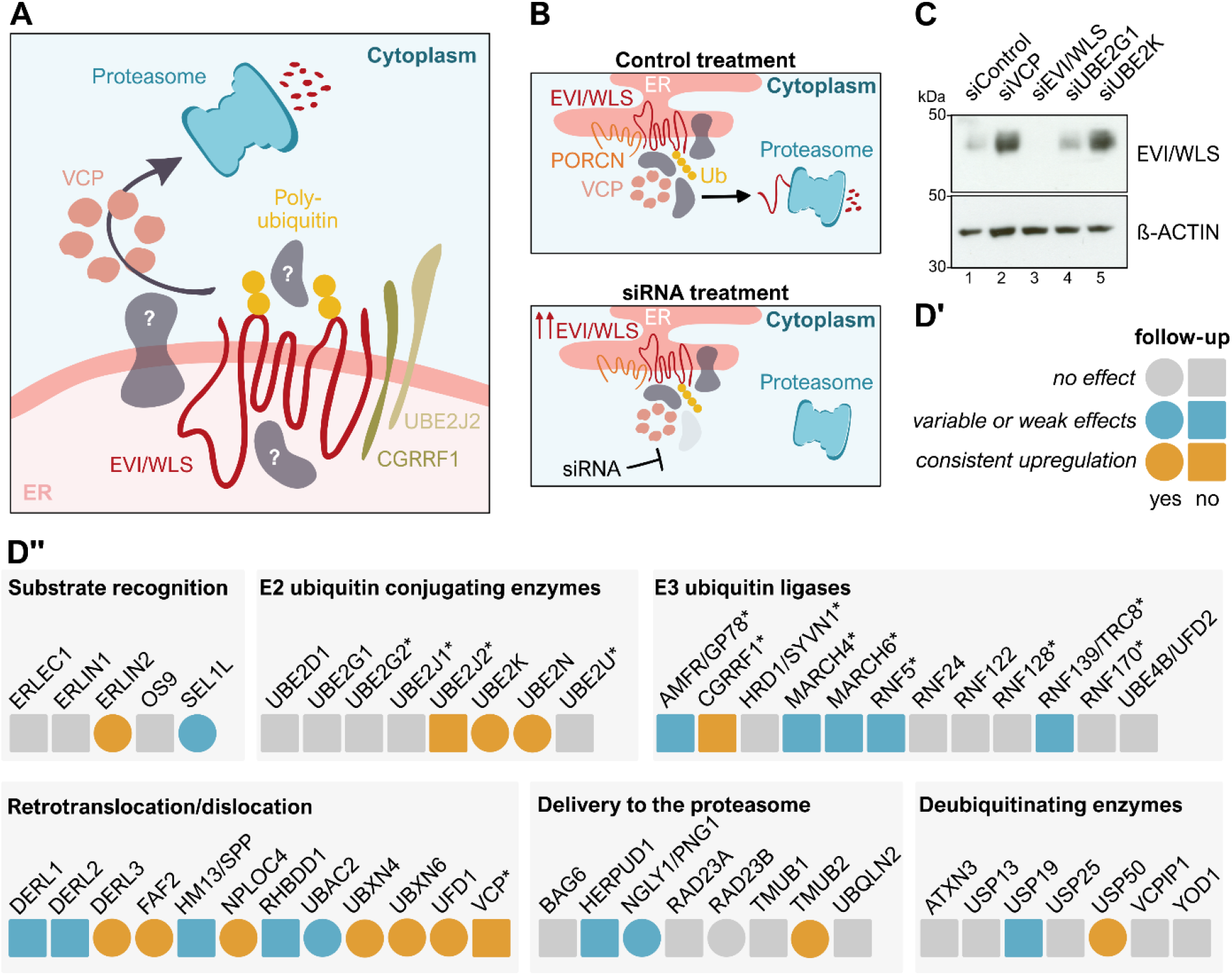
siRNA-based mini-screen identifies novel candidates involved in the degradation of EVI/WLS. **A**. Schematic representation of the current understanding of the ERAD of EVI/WLS and open questions. EVI/WLS is ubiquitinated by CGRRF1 and UBE2J2 before being extracted from the ER membrane with the help of VCP and degradation by the proteasome. **B**. Schematic illustration of the principle behind the screening procedure with siRNAs. EVI/WLS protein accumulates if the siRNA targets a mRNA encoding a protein important for EVI/WLS degradation. ER = endoplasmic reticulum, Ub = poly-ubiquitin chain **C**. EVI/WLS protein levels were analysed after siRNA mediated knock-down of target genes. Increased EVI/WLS protein levels compared to siControl treatment indicated the possible involvement of the candidate in EVI/WLS’s ERAD processing. HEK293T cells were treated with the indicated siRNAs for 72 h. β-ACTIN served as loading control. Western blots are representative of three independent experiments. kDa = kilodalton **D**. Results of the siRNA-based mini-screen represented as heat-map. Candidates without effect are marked in grey, candidates with variable or weak effects are marked in blue and candidates that show a strong and consistent upregulation of EVI/WLS are marked in yellow. Circles indicate follow-up. Asterisks indicate genes that were previously tested in Glaeser et al., 2018. A detailed table including gene accession numbers and phenotypes in HEK293T and A375 cells can be found in Appendix Table S1, the Western blots underlying this analysis are shown in Figure S1.

In this study, we performed a focused RNAi- and Western blot-based screen on EVI/WLS protein abundance to identify factors regulating its stability. We found that its degradation is initiated by interaction with ERLIN2, which precedes EVI/WLS ubiquitination. Thereafter, ubiquitinated EVI/WLS is linked to VCP by its interaction with FAF2 and UBXN4. Furthermore, we demonstrate that human EVI/WLS is additionally ubiquitinated by UBE2N and UBE2K (beside UBE2J2) and modified with ubiquitin of various linkage types (K11, K48, and K63) which has important consequences for its protein stability and function. Using melanoma cells which express high levels of WNT ligands, we show that endogenous EVI/WLS proteins are ubiquitinated and degraded even in the presence of WNTs. Thus, our data provides important insights into the mechanism of the recognition and degradation of EVI/WLS, an endogenous mammalian substrate of regulatory ERAD, and further emphasises the link between ubiquitination and WNT signalling.

## RESULTS

### A focused screen for ERAD candidate genes

Several proteins are involved in the recognition and retrotranslocation/dislocation of ERAD substrates, but additional regulators of EVI/WLS remain elusive. To address this in a systematic manner, we performed a focused siRNA and Western blot-based screen in HEK293T cells. As read-out we used the EVI/WLS protein level after treatment with a pool of four siRNAs and increased protein levels indicated impaired degradation (Figure 1B). In summary, we tested 52 candidates, including those previously tested by Gläser et al., 2018 (Figures 1C, 1D, S1 and Suppl Table S1).

In the follow-up experiments, we investigated 15 selected candidates in more detail by assessing the effects of the respective single siRNAs on EVI/WLS protein and, in selected cases, on mRNA level. RT-qPCR was used to analyse knock-down efficiencies and to exclude transcriptional regulation of EVI/WLS, thus highlighting post-translational effects. siEVI/WLS was used as on-target control and siVCP as positive control. siVCP has been previously shown to increase endogenous EVI/WLS protein levels without affecting EVI/WLS mRNA levels (Gläser et al., 2018). The silencing of mRNA and protein expression by siEVI/WLS and siVCP were efficient (Figures S1, S2 and S3) and VCP downregulation induced upregulation of EVI/WLS protein levels, as described previously. The majority of the tested siRNAs efficiently reduced mRNA levels to values between 5 % and 20 % of the control without affecting EVI/WLS protein expression. However, 10 candidates were not followed up due to variation in the Western blot experiments between different single siRNAs and biological replicates indicating non-target effects of the siRNAs (Figures S2 and S3).

### EVI/WLS protein levels are regulated by our candidate proteins

Five genes from our screen (*FAF2, ERLIN2, UBXN4*, and the E2 ubiquitin conjugating enzymes *UBE2K* and *UBE2N*) were selected for further analysis due to their consistent up-regulation of EVI/WLS protein levels without changes in *EVI/WLS* mRNA levels (Figures 2, 4A,B).

**Figure 2.**
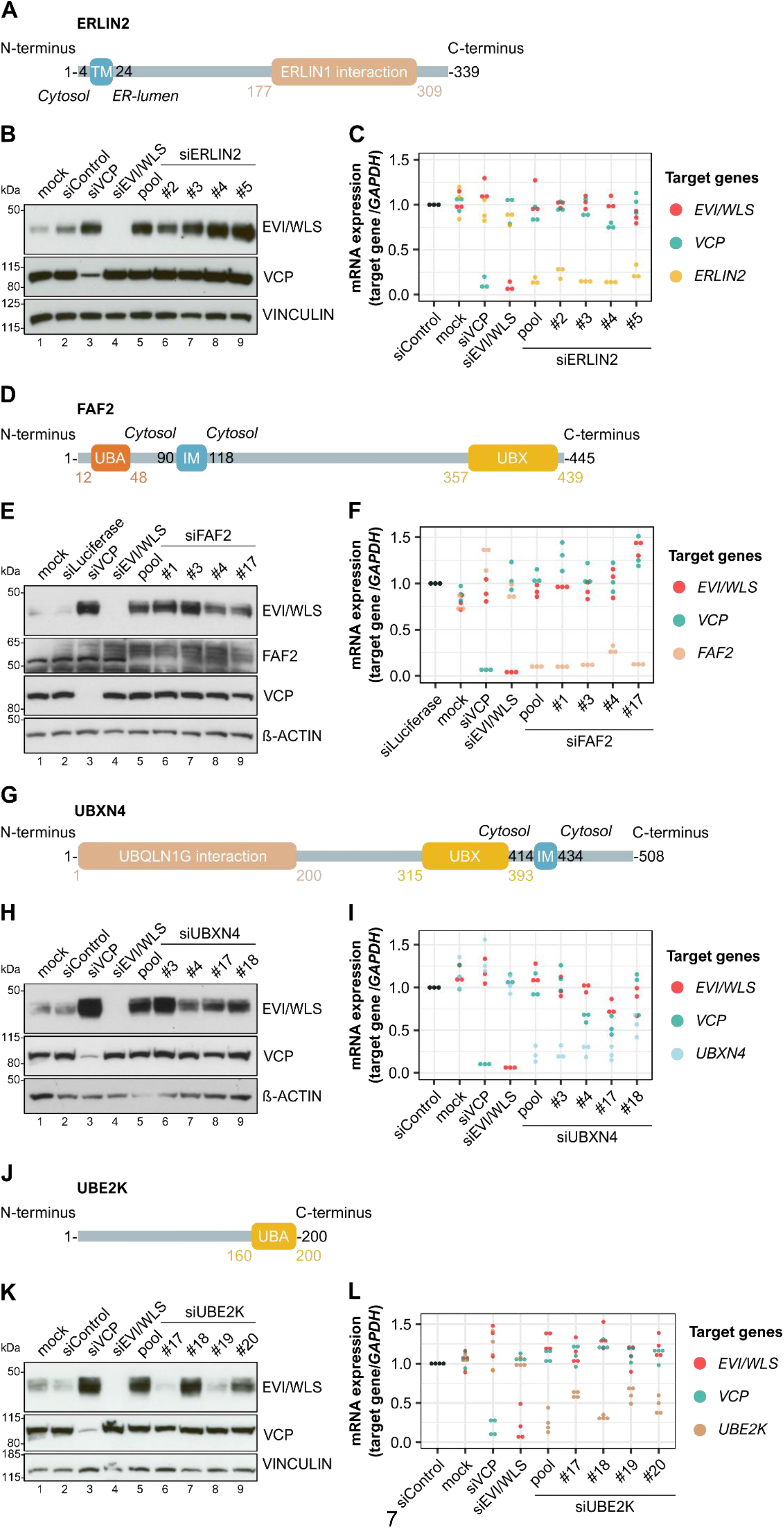
ERLIN2, FAF2, UBXN4, and UBE2K regulate endogenous EVI/WLS on protein level. **A, D, G, J**. Schematic representations of the proteins ERLIN2, FAF2, UBXN4, and UBE2K according to UniProt IDs: O94905, Q96CS3, Q92575, and P61086 respectively. Numbers indicate amino acid positions. TM = transmembrane domain, IM = intramembrane domain, UBA = ubiquitin associated domain important for binding to ubiquitin, UBX = ubiquitin regulatory X domain important for binding to VCP, ER = endoplasmic reticulum **B, C, E, F, H, I, K, L**. Knock-down of ERLIN2, FAF2, UBXN4, or UBE2K increased EVI/WLS protein levels but had no effect on *EVI/WLS* mRNA expression. HEK293T cells were treated with the indicated siRNAs for 72 h. siRNAs targeting *ERLIN2, FAF2*, or *UBXN4* were used as either single siRNAs or an equimolecular mix of all four respective siRNAs (pool). siRNA numbers are according to the manufacturer. **B, E, H, K**, VINCULIN or β-ACTIN served as loading control. Western blots are representative of three independent experiments. kDa = kilodalton. **C, F, I, L**, Target gene expression was normalised to siControl treatment and *GAPDH* served as reference gene. Individual data points of three or four independent experiments are shown.

To test whether these proteins bind to EVI/WLS, we performed immunoprecipitation studies. Here, we generated FLAG-tagged overexpression constructs of ERLIN2, FAF2, UBXN4, and UBE2K to investigate their interaction with endogenous EVI/WLS and VCP. PORCN-FLAG was used as a positive control since it has been previously shown that it binds to EVI/WLS and VCP (Glaeser et al., 2018). Indeed, our immunoprecipitation studies demonstrate that endogenous EVI/WLS interacts with ERLIN2-FLAG, FAF2-FLAG, and UBXN4-FLAG (Figure 3A,C,D, respectively). Furthermore, we detected an interaction between endogenous ERLIN2 and FAF2 with PORCN-FLAG (Figure S4A,B). We could also confirm previously described interactions within the ERAD machinery, such as between FAF2 and ERLIN2 (Figure S4A,B, Christianson et al., 2012), and between VCP and the UBX domain containing proteins FAF2 and UBXN4 (Figure 3C,D, Schuberth & Buchberger, 2008), but interestingly not between FAF2 and UBXN4 (Figure S4A). ERLIN1 is an important interaction partner of ERLIN2, but it did not influence EVI/WLS protein levels in our screen (Figures 1D and S1). Accordingly, our pull-down experiments revealed that it interacts with endogenous ERLIN2, but not with EVI/WLS or VCP (Figures 3B, S4B).

**Figure 3.**
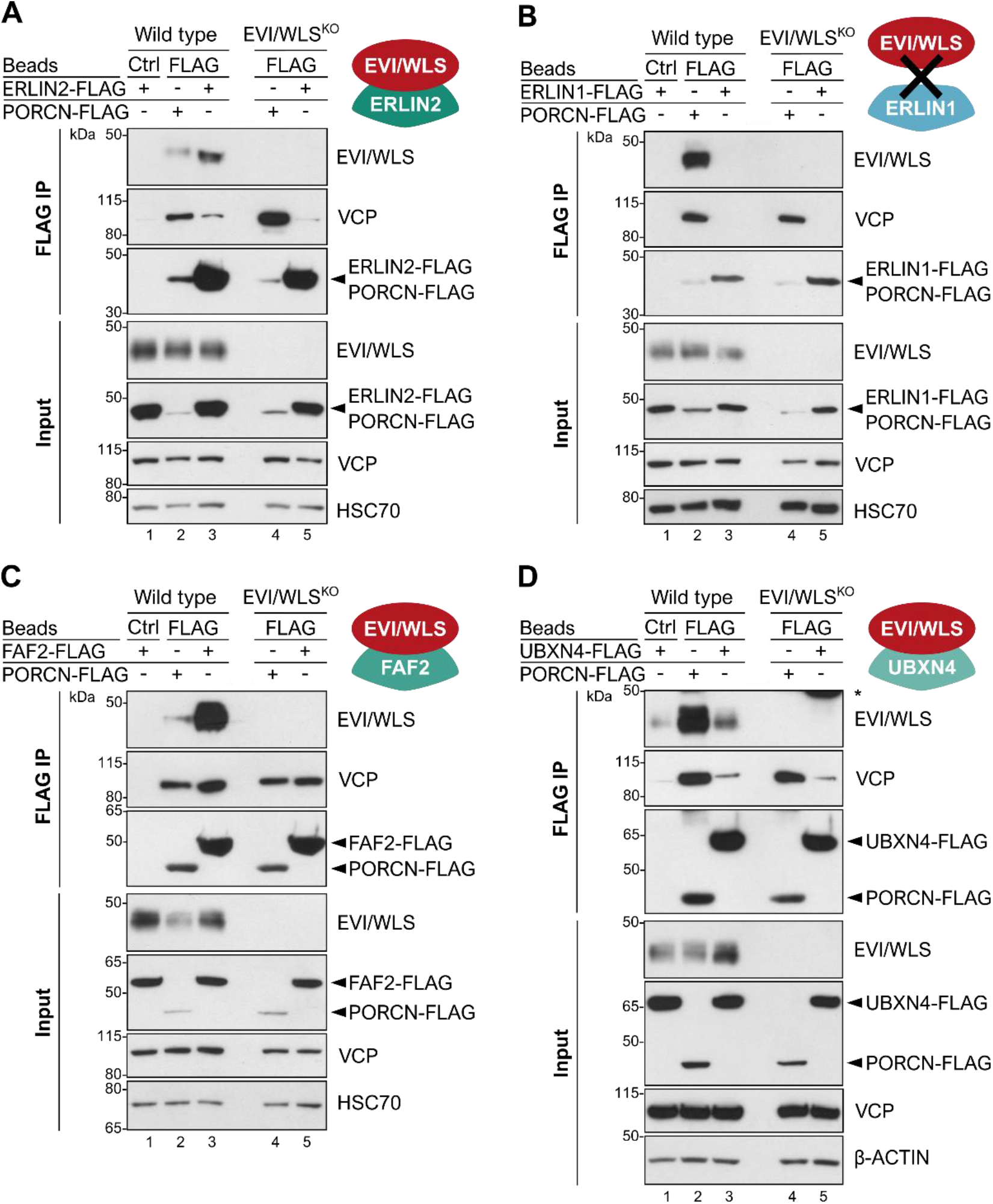
ERLIN2-FLAG, FAF2-FLAG, and UBXN4-FLAG interact with endogenous EVI/WLS. Immunoprecipitation (IP) experiments confirmed interaction between endogenous EVI/WLS and ERLIN2-FLAG (**A**), FAF2-FLAG (**C**), and UBXN4-FLAG (**D**) but not ERLIN1-FLAG (**B**). HEK293T wild type and EVI/WLS knock-out (EVI/WLS^KO^) cells were transfected with ERLIN1-FLAG, ERLIN2-FLAG, FAF2-FLAG, UBXN4-FLAG, or PORCN-FLAG overexpression plasmids. After 48 h, total cell lysates were sampled for input control or used for FLAG IP to precipitate FLAG-tagged proteins and their interaction partners. HSC70 or β-ACTIN served as loading control. Western blots are representative of three independent experiments. KDa = kilodalton

Our screen also identified two E2-conjugating enzymes, UBE2K and UBE2N, which regulate EVI/WLS protein levels. However, we were unable to detect an interaction between FLAG-UBE2K and EVI/WLS in immunoprecipitation experiments (Figure S4C), presumably due to the transient nature of the interaction and the stringent pull-down conditions used in this study.

RNAi-induced knock-down of UBE2N increased EVI/WLS protein levels in HEK293T cells proportional to siRNA efficiency (Figures 4A,B). UBE2N (UBC13) is an E2 enzyme with K63 linkage specificity required for protein localisation and endosomal trafficking (Akutsu et al., 2016; Erpapazoglou et al., 2014; Swatek & Komander, 2016), processes which are also essential for the function of EVI/WLS. In addition to UBE2N, we also tested the effect of its catalytically inactive interaction partners UBE2V1 and UBE2V2 (Andersen et al., 2005; McKenna et al., 2001) on EVI/WLS protein levels (Figure 4). Importantly, knock-down of *UBE2N* and its interaction partners not only increased EVI/WLS protein levels, but also the secretion of WNT ligands in HEK293T cells upon WNT3 or NanoLuciferase-WNT3 overexpression, indicating a possible modulatory effect on WNT signalling in general (Figure 4D,E). Indeed, Zhang et al. described the importance of Ubc13/UBE2N for WNT-dependent processes in worms (Zhang et al., 2018). The effects of UBE2V1 knock-down on protein stability and WNT secretion were more variable and suggested indirect effects, maybe *via* stability of the UBE2N complex (Figure 4D,E). Under these conditions, secretion of WNT3 was not increased after knock-down of VCP (Figure 4D,E).

**Figure 4.**
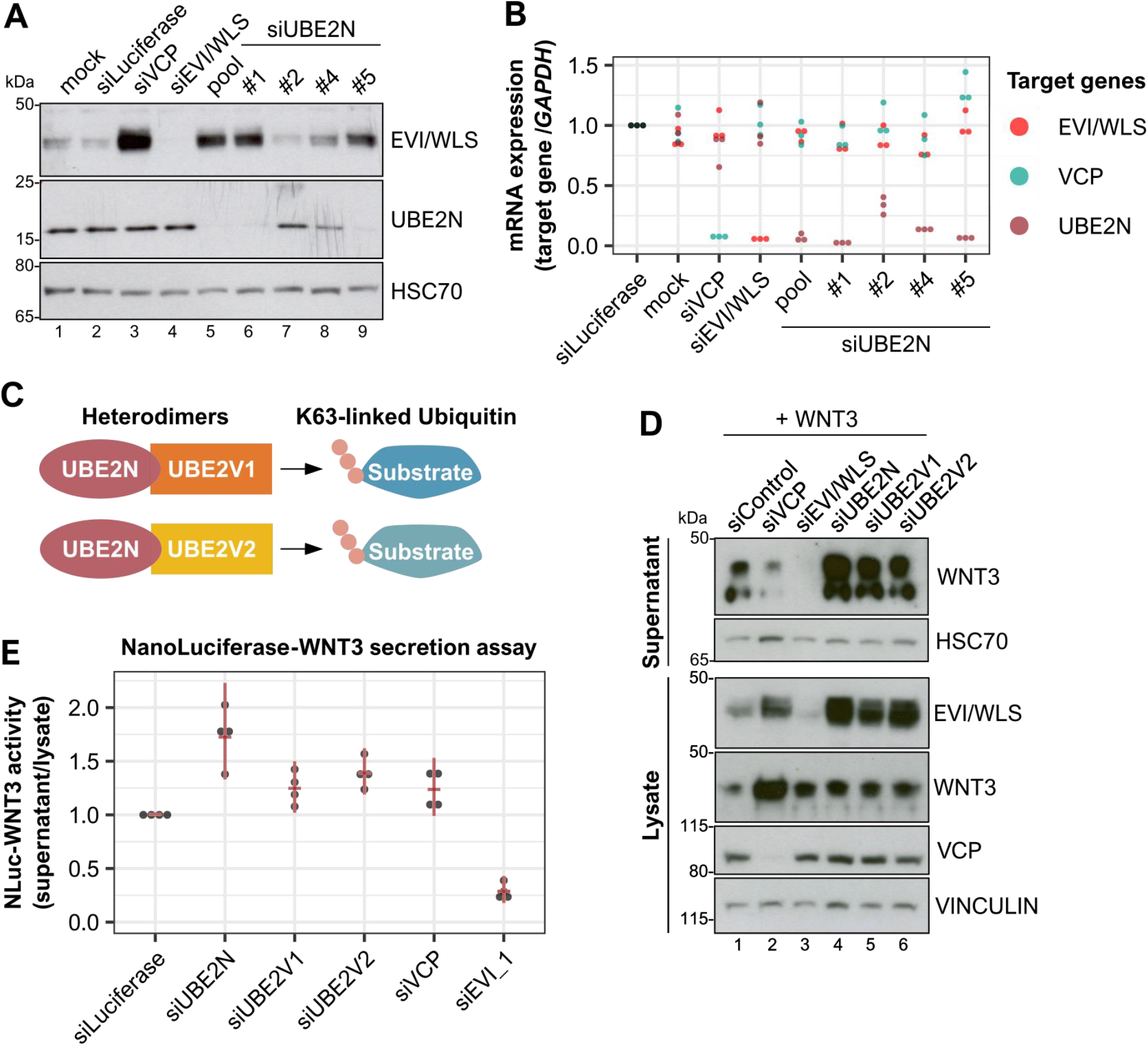
UBE2N and UBE2V1/UBE2V2 regulate EVI/WLS protein levels and WNT secretion. **A**. Knock-down of UBE2N increased EVI/WLS protein levels. HEK293T cells were treated with indicated siRNAs for 72 h. siRNAs targeting *UBE2N* were used as either single siRNAs or an equimolecular mix of all four respective siRNAs (pool). HSC70 served as loading control. **B**. mRNA expression analyses showed mostly potent gene silencing by pooled or single siRNAs with little effects on other investigated mRNAs. HEK293T cells were treated with the indicated siRNAs for 72 h. Each gene’s mRNA was targeted by either single siRNAs or an equimolecular mix of all four respective siRNAs (pool) to analyse their effect on mRNA expression. Target gene expression was normalised to siLuciferase treatment and *GAPDH* served as reference gene. Individual data points from three independent experiments are shown. **C**. Schematic representation of the formation of active complexes by UBE2N with UBE2V1 or UBE2V2 to modify target substrates with K63-linked ubiquitin. **D**. Knock-down of UBE2N, UBE2V1, and UBE2V2 in combination with WNT3 overexpression increased EVI/WLS protein levels and WNT secretion compared to control treatment. HEK293T cells were treated with indicated siRNAs. 24 h after siRNA transfection, cells were additionally transfected with WNT3 plasmid. VINCULIN or HSC70 served as loading control. **E**. NanoLuciferase (NLuc)-WNT3 secretion was elevated after knock-down of UBE2N. HEK293T cells were treated with the indicated siRNAs and transfected with NLuc-WNT3 and Firefly Luciferase 24 h later. 48 h later, NLuc activity was determined in the cells’ supernatant and normalised to NLuc and Firefly activity in the cell lysates. Data points derived from four independent experiments with mean and confidence intervals are shown. Western blots are representative of three independent experiments. kDa = kilodalton

### EVI/WLS is ubiquitinated and degraded in the presence of endogenous WNT ligands

HEK293T cells are commonly used as a model to analyse WNT signalling due to their low endogenous WNT secretion. The striking observation that ubiquitination of EVI/WLS might influence WNT ligand secretion indicates that it might regulate WNT signalling itself. Hence, we were intrigued by the ubiquitination status of EVI/WLS in cells with high endogenous WNT ligand production. Melanoma is a skin cancer with poor prognosis in advanced stages (Schadendor et al., 2018) and tumour progression and metastasis are associated with the expression of non-canonical WNT ligands (Webster et al., 2015). For this reason, we chose the melanoma cell line A375 for further studies, which has been shown to express high amounts of WNT proteins, most notably WNT5A (Yang et al., 2012). Since the overexpression of WNT ligands in HEK293T cells led to the stabilisation of EVI/WLS protein (Glaeser et al., 2018), we wanted to investigate whether the effect can be reversed in A375 cells by using LGK974, an inhibitor of the acyl-transferase Porcupine (PORCN), thus preventing the secretion of WNT ligands. As expected, LGK974 treatment abolished the secretion of WNT5A, and interestingly reduced EVI/WLS protein levels in the cell lysates without affecting EVI/WLS mRNA expression (Figures 5A,B,C, note that the antibody used in these studies recognises both WNT5A and WNT5B).

**Figure 5.**
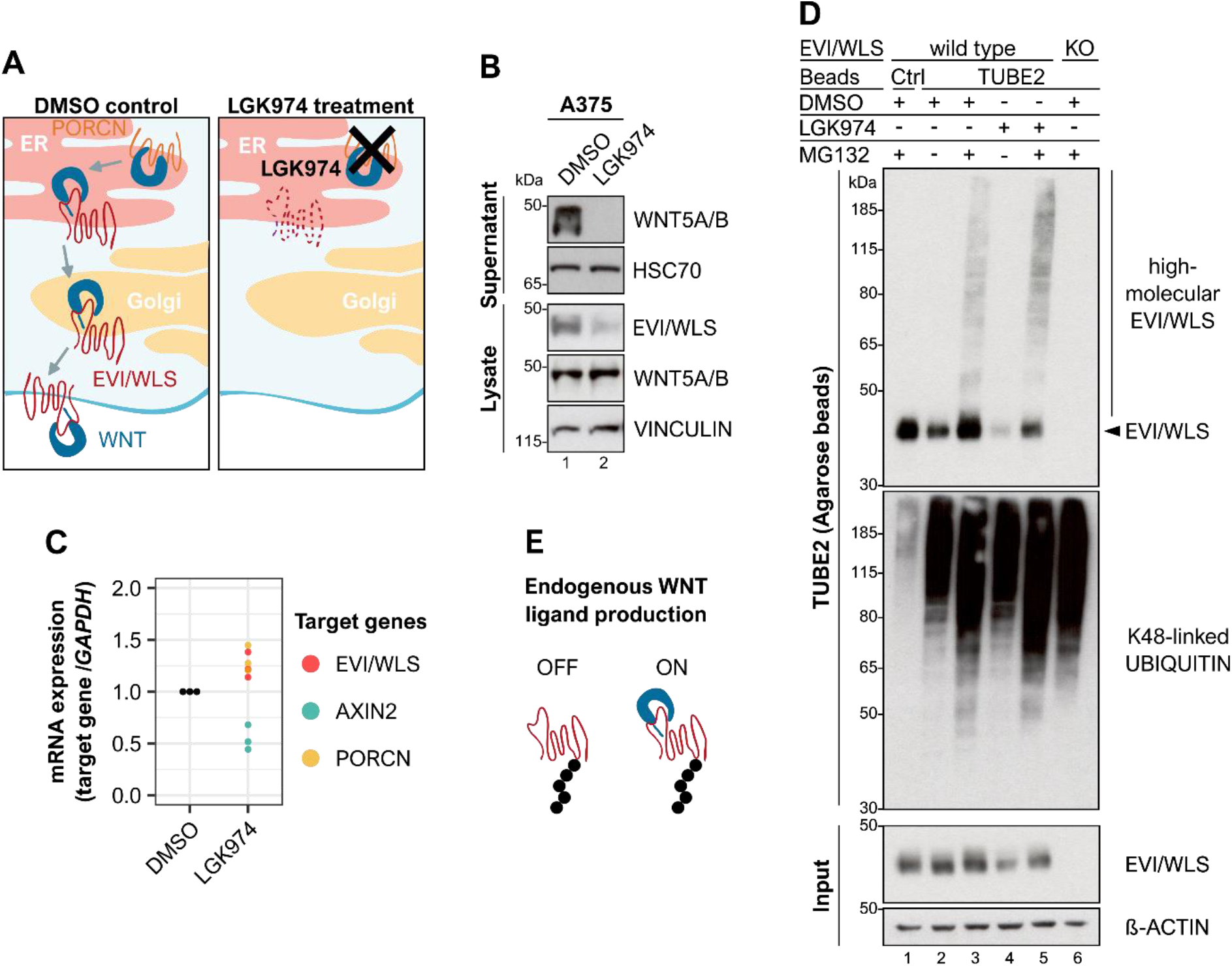
EVI/WLS is ubiquitinated in cells with endogenous WNT ligands. **A**. Schematic illustration of LGK974’s mode of action. LGK974 prevents WNT ligands from being lipid-modified in the endoplasmic reticulum (ER) by inhibiting the acyl-transferase PORCN. Un-lipidated WNTs cannot associate with EVI/WLS and are not secreted from the WNT producing cell. **B, C**. LGK974 treatment reduced intracellular EVI/WLS levels and abolished the secretion of WNT5A/B ligands, without reducing EVI/WLS gene expression. A375 melanoma cells were treated with LGK974 (10 µM) for 96 h with daily medium changes. **B**, secreted proteins were precipitated from the supernatant using Blue Sepharose VINCULIN or HSC70 served as loading control. **C**, Target gene expression was normalised to DMSO treatment and *GAPDH* served as reference gene. Individual data points from three independent experiments are shown. **D**. Ubiquitinated EVI/WLS accumulated after inhibition of the proteasome independent of LGK974 treatment. A375 melanoma wild type and EVI/WLS knock-out (EVI/WLS^KO^) cells were treated with LGK974 (10 µM) for 96 h with daily medium changes. 24 h before harvest, samples were treated with the proteasome inhibitor MG132 (1 µM) as indicated. Then, total cell lysates were sampled for input control or used for TUBE2 (agarose) pull-down to precipitate poly-ubiquitinated proteins. Ubiquitin non-binding Control (Ctrl) Agarose Beads showed level of unspecific binding and EVI/WLS^KO^ cells confirmed specificity for EVI/WLS. β-ACTIN served as loading control. TUBE = tandem ubiquitin binding entity **E**. EVI/WLS is ubiquitinated in the presence and absence of endogenous WNT ligands Western blots are representative of three independent experiments. kDa = kilodalton

Next, we were interested in the ubiquitination status of EVI/WLS on an endogenous level. For this reason, we used Tandem Ubiquitin Binding Entities (TUBEs) to enrich for ubiquitinated proteins followed by Western blot analysis. Combinatorial treatment with LGK974 and the proteasome inhibitor MG132 increased the accumulation of high molecular weight bands of EVI/WLS, as expected. Surprisingly, treatment with MG132 and DMSO as vehicle control for LGK974 resulted in the accumulation of ubiquitinated EVI/WLS as well, indicating EVI/WLS is ubiquitinated in the absence and presence of lipid-modified endogenous WNT ligands (Figure 5D,E). These results show that EVI/WLS protein levels depend on the availability of mature WNT ligands in cells with high endogenous WNT signalling, but a certain fraction of the EVI/WLS protein is ubiquitinated and targeted for degradation even in the presence of WNT ligands. To identify genes that mediate the degradation of EVI/WLS in A375 cells, we silenced candidate genes by RNAi in the presence of LGK974 and analysed EVI/WLS protein levels by Western blot. Among the tested genes, only the knock-down of UBE2J2, CGRRF1, and VCP consistently elevated EVI/WLS protein levels, in line with our previous results in HEK293T cells (Figure S5 A,B,C,D, Glaeser et al., 2018).

### ERLIN2 connects EVI/WLS to the ubiquitination machinery

It is known that EVI/WLS is ubiquitinated by UBE2J2 and CGRRF1 (Fenech et al., 2020; Glaeser et al., 2018) and that the UBE2J2 ortholog in yeast (Ubc6) is indispensable for priming a broad range of substrates with monoubiquitin or K11-linked ubiquitin dimers (Weber et al., 2016; Xu et al., 2009). However, so far the kind of ubiquitin linkage types present on EVI/WLS and the sites of its modification have not been identified, since the TUBEs used to detect ubiquitination in Figure 5D were non-selective and unable to distinguish between different ubiquitin linkage types. Therefore, we used mutant ubiquitin constructs which only allow one linkage type, either K11, K48, or K63 (Clague et al., 2015; Tsuchiya et al., 2018; Xu et al., 2009). Intriguingly, we detected ubiquitinated EVI/WLS bands upon wild type, K11, K48, or K63 ubiquitin-HA overexpression, but not in control conditions (Figure S5E), indicating the presence of multiple linkage types. This data also supports the hypothesis that EVI/WLS is modified by multiple E2 enzymes with different linkage type specificity.

Our results so far showed that ERLIN2, FAF2, UBXN4, UBE2K, and UBE2N could regulate EVI/WLS protein levels and that EVI/WLS is stabilised in cells with endogenously active WNT signalling but can still be ubiquitinated and degraded. However, it remained elusive how these candidates can influence the ubiquitination of EVI/WLS. We hypothesised that UBE2K and UBE2N would modify EVI/WLS with K48- and K63-linked ubiquitin, respectively (Figure S5E).

Linkage-type specific TUBEs can detect differences in K48- or K63-linked ubiquitination, especially, if the substrate protein is modified in parallel with several poly- or mono-ubiquitins and differences in over-all ubiquitination are small. UBE2K can synthesise K48-linked ubiquitin chains (Z. Chen & Pickart, 1990; Middleton & Day, 2015) and accordingly, knock-down of UBE2K resulted in reduced K48-ubiquitinated EVI/WLS (Figure 6A). Similarly, silencing of UBE2N and UBE2V2 strongly reduced K63-specific endogenous high-molecular EVI/WLS bands (Figure 6B). This data indicates that UBE2K mediates K48-linked ubiquitination of EVI/WLS whereas UBE2N and UBE2V2 mediate K63-linked ubiquitination of EVI/WLS in human cells. While K63-linked ubiquitin chains can direct substrates towards lysosomal degradation (R.-H. Chen et al., 2019; Pohl & Dikic, 2019), the accumulation of K63-ubiquitination of EVI/WLS after MG132 treatment indicates proteasomal degradation.

**Figure 6.**
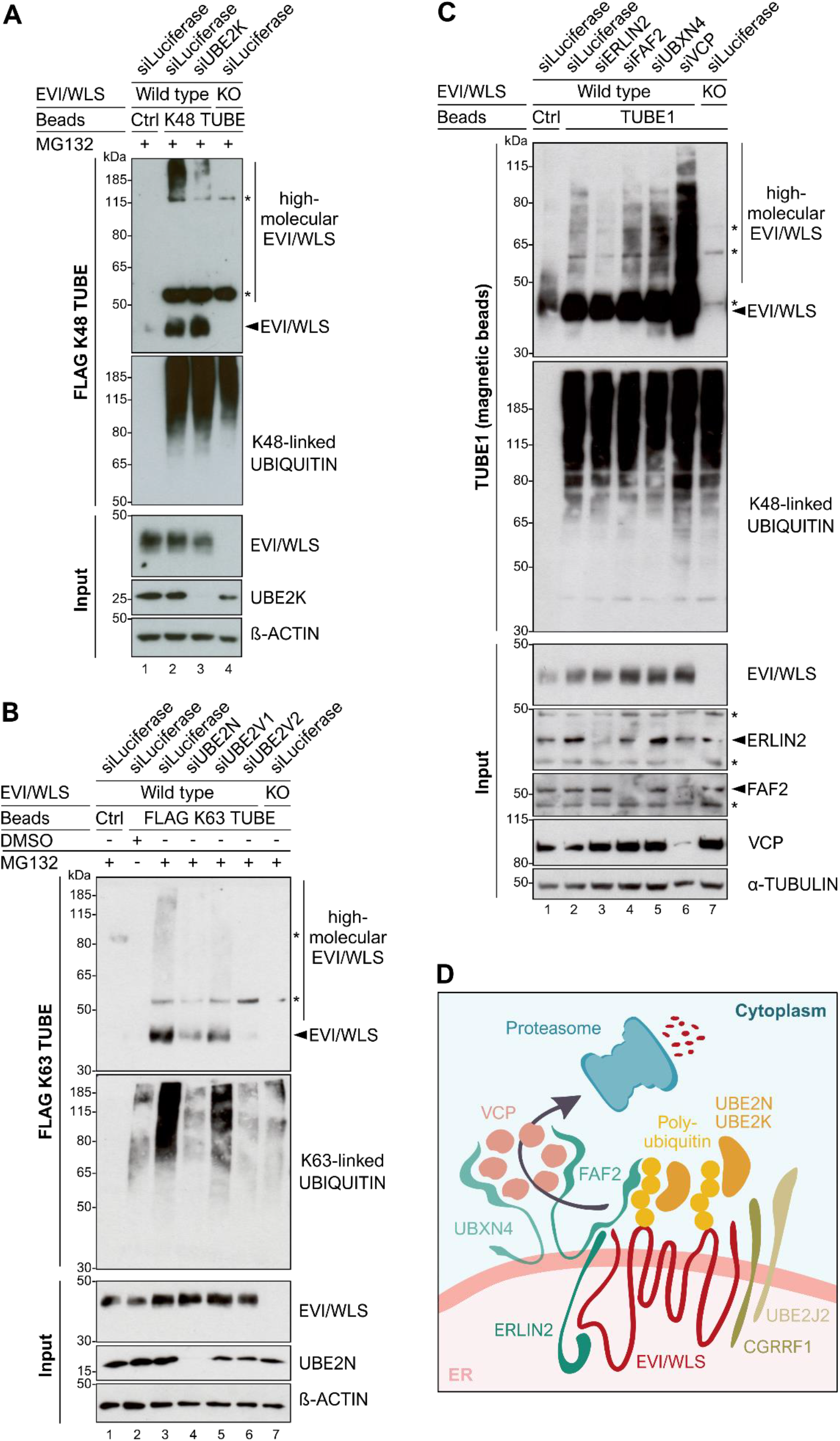
ERLIN2 links EVI/WLS to the ubiquitination machinery. **A**. FLAG K48 specific TUBE pull-down confirmed that EVI/WLS is modified with K48-linked ubiquitin chains by UBE2K. A375 melanoma wild type and EVI/WLS knock-out (EVI/WLS^KO^) cells were treated with indicated siRNAs for 72 h. 24 h before harvest, samples were treated with the proteasome inhibitor MG132 (1 µM). Total cell lysates were sampled for input control or used for FLAG K48 TUBE pull-down to specifically precipitate proteins modified with K48-linked poly-ubiquitin. β-ACTIN served as loading control. **B**. FLAG K63 specific TUBE pull-down confirmed that EVI/WLS is modified with K63-linked ubiquitin chains by UBE2N. A375 melanoma wild type and EVI/WLS knock-out (EVI/WLS^KO^) cells were treated with indicated siRNAs for 72 h. 24 h before harvest, samples were treated with the proteasome inhibitor MG132 (1 µM). Total cell lysates were sampled for input control or used for FLAG K63 TUBE pull-down to specifically precipitate proteins modified with K63-linked poly-ubiquitin. β-ACTIN served as loading control. **C**. Knock-down of ERLIN2 reduced the ubiquitination of EVI/WLS, while knock-down of FAF2 and UBXN4 increased it. A375 wild type and EVI/WLS knock-out (KO) cells were treated with the indicated siRNAs for 72 h. Then, total cell lysates were sampled for input control or used for TUBE1 pull-down to precipitate poly-ubiquitinated proteins. α-TUBULIN served as loading control. **D**. Schematic representation of the ERAD of EVI/WLS. EVI/WLS is recognised by ERLIN2 and ubiquitinated by CGRRF1 and UBE2J2 as well as UBE2K and UBE2N. The latter adds K63-linked ubiquitin chains to EVI/WLS. Ubiquitinated EVI/WLS binds FAF2 and UBXN4, which recruit VCP to the ER membrane and mediate the delivery of EVI/WLS to the proteasome. Asterisks mark nonspecific signals. Western blots are representative of three independent experiments. kDa = kilodalton, TUBE = tandem ubiquitin binding entity

To determine whether ERLIN2, FAF2 and UBXN4 are also required for EVI/WLS ubiquitination, we performed precipitations using pan-ubiquitin specific TUBE1 magnetic beads in combination with RNAi mediated knock-down of these candidate genes. Silencing of ERLIN2 led to a reduction of high-molecular EVI/WLS bands, whereas knock-down of FAF2 and UBXN4 resulted in an increase of polyubiquitinated EVI/WLS (Figure 6C). Together, this data suggests that ERLIN2 functions as a bridge connecting EVI/WLS to the ubiquitination machinery and that FAF2 and UBXN4 interact with EVI/WLS after it is ubiquitinated, but before it is delivered to the proteasome. We used siVCP as a positive control, which actually showed the strongest accumulation of high-molecular Evi/Wls bands compared to all conditions (Figure 6C), presumably indicating that EVI/WLS is subject to several parallel recruitment mechanisms which all culminate in dislocation by VCP.

Accordingly, we envision the following model how EVI/WLS is ubiquitinated and subjected to ERAD: ERLIN2 serves as an important link between EVI/WLS and other ERAD components and potentially helps to recruit the ubiquitination machinery, consistent at least of UBE2K, UBE2N, UBE2J2, and CGRRF1. Poly-ubiquitinated EVI/WLS interacts with FAF2 and UBXN4, which recruit VCP to the ER membrane, resulting in the dislocation and eventually the degradation of EVI/WLS (Figure 6D).

## DISCUSSION

Cellular signalling pathways frequently regulate and are regulated by protein stability. In this study, we investigated the ubiquitination of the WNT ligand cargo protein EVI/WLS and how it is linked to the ERAD machinery. Hence, we performed a small-scale loss-of function screen using EVI/WLS protein stability as a phenotypic readout (Figure 1). By this means, we identified five candidates which increased EVI/WLS protein levels upon knock-down: ERLIN2, FAF2, UBXN4, UBE2K, and UBE2N (Figures 1,2,4). FAF2 and UBXN4 contain VCP interaction domains and are anchored at the ER membrane by an ‘intramembrane’ domain, which leaves both their N- and C termini facing the cytoplasm (Liang et al., 2006; Meyer & Weihl, 2014; Mueller et al., 2008; Schuberth & Buchberger, 2008). This allows them to hold a firm grip on VCP and to support it during the generation of mechanical force by ATP dependent protein extraction from the ER (Hirsch et al., 2009). Our data indicates an interaction between EVI/WLS and ERLIN2 already prior to ubiquitination (Figure 6C), suggesting a role of ERLIN2 as linker between EVI/WLS and the ERAD machinery, similar as described for HMG-CoA reductase and IP3R (Jo et al., 2011; Pearce et al., 2007, 2009; Y. Wang et al., 2009). Immunoprecipitation experiments confirmed the interaction of FAF2 and ERLIN2, which was reported in a previous study by mapping ERAD component interactions (Christianson et al., 2012). The additional interaction with VCP and PORCN indicates either the formation of a large complex at the ER membrane prior to EVI/WLS degradation or a sequential recruitment of these proteins.

It has been described previously that the complex of ERLIN1 and ERLIN2 is important for the degradation of IP3R (Pearce et al., 2009), however we found that ERLIN1 did not regulate EVI/WLS protein levels and did not interact with EVI/WLS (Figures 1,3). This is in line with the described ERLIN1-independent role of ERLIN2 for the degradation of HMG-CoA reductase (Jo et al., 2011), indicating several parallel mechanisms involving ERLIN2 that probably depend on additional interaction partners. It is assumed that most ERAD related proteins have been identified in yeast and mammals (Christianson & Ye, 2014). However, it cannot be excluded that there are additional ERAD-associated proteins that regulate EVI/WLS protein levels which could not be detected in our assay due to non-specific siRNAs or cell line dependency.

We found that endogenous EVI/WLS is modified with ubiquitin of several linkage types (K11, K48, and K63, Figures 6,S5E), as was recently described for other ERAD clients as well (Leto et al., 2019). The presence of K63-linked ubiquitin chains indicate a role in EVI/WLS endocytosis and trafficking as well as degradation. It is well known that EVI/WLS associates with WNT ligands in the ER of WNT secreting cells and helps to shuttle them to the cell surface (Bänziger et al., 2006; Bartscherer et al., 2006; Goodman et al., 2006; Yu et al., 2014). Afterwards, EVI/WLS is endocytosed with the help of CLATHRIN and recycled back to Golgi and ER in a retromer dependent process, where it can bind again to WNTs (Belenkaya et al., 2008; Port et al., 2008). Upon inhibition of EVI/WLS trafficking, it is shuttled to the lysosomes for degradation (Franch-Marro et al., 2008; Gross et al., 2012; Yang et al., 2008). Recently, a study in *Caenorhabditis elegans* found that the knock-out of UBC13 (human UBE2N, E2 ubiquitin conjugating enzyme with K63 specificity) also disrupted MIG-14/EVI/WLS trafficking and diverted it to lysosomes whereas they did not show direct ubiquitination of MIG-14/EVI/WLS (Zhang et al., 2018). In agreement with this study, we show here that human EVI/WLS is modified with K63-linked ubiquitin by UBE2N and UBE2V2 (Figure 6B). Moreover, loss of UBE2N and UBE2V2 led to increased EVI/WLS protein levels and WNT secretion (Figure 4). The apparent reduction in secretion of WNT3 after knock-down of VCP could be due to its viability effect or because WNT3 might be retained in the secretory machinery so that it cannot be secreted. Notably, VCP has pleiotropic effects and different phenotypes, including cell death, can be observed according to the experimental conditions, such as e.g. timing and length (Glaeser et al., 2018).

We did not yet identify the E3 ubiquitin ligase which is required for UBE2N-mediated ubiquitination of EVI/WLS, but our data and recently published high-throughput studies indicate that it is presumably not an ER-membrane associated protein, but rather cytosolic (Fenech et al., 2020; Leto et al., 2019; Glaeser et al., 2018). It should also be considered that earlier *in vitro* and structural studies showed that UBE2N∼ubiquitin together with UBE2V2 can adopt its active conformation even in the absence of an E3 ligase, suggesting E3 independent ubiquitin chain elongation (McKenna et al., 2001; Pruneda et al., 2011). However, sophisticated real-time FRET analysis did not reveal ubiquitin transfer events in the absence of an E3 (Branigan et al., 2020).

Our data indicate that EVI/WLS is additionally ubiquitinated by UBE2K (Figure 6A), in addition to the previously described UBE2J2 and CGRRF1 (Glaeser et al., 2018). The yeast homolog of UBE2J2 (Ubc6) was reported to prime substrates with short K11-linked ubiquitin modifications, which are presumably not sufficient to recruit the degradation machinery (Mehrtash & Hochstrasser, 2019; Tsuchiya et al., 2018; Weber et al., 2016; Xu et al., 2009). Therefore, it is tempting to speculate that UBE2K would elongate these initial modifications by UBE2J2 in mammalian cells and allow successful interaction with downstream factors, potentially even without an associated E3 enzyme (Middleton & Day, 2015; Rodrigo-Brenni & Morgan, 2007). It is currently still unclear which amino acids of EVI/WLS are ubiquitinated. Publicly available mass spectrometry data report ubiquitination at several lysine residues across its protein sequence. However, UBE2J2/Ubc6 can also modify hydroxylated amino acids (Weber et al., 2016), such as serine and threonine, which are not detected with standard mass spectrometry approaches. Therefore, further studies are required to characterise the composition and localisation of ubiquitin modifications carried by EVI/WLS.

The presented data provides insights into the degradation mechanism of an endogenous substrate of mammalian regulatory ERAD, which seems to function independently of the well-studied E3s HRD1, GP78, and MARCH6 and engages with various E2 ubiquitin conjugating enzymes. Surprisingly, the knock-down of UBE2G2 had no effect on EVI/WLS protein abundance (Figure 1), although UBE2G2 was recently reported to be required for the degradation of several other ERAD clients (Leto et al., 2019). It will be important to test in the future whether other substrates of regulatory ERAD are also UBE2G2 independent or if this is a cell type or assay specific effect.

There are still open questions regarding EVI/WLS degradation, and especially the nature of a potential ER membrane channel protein for its dislocation remains elusive, as our screen was not able to identify such a protein. While the extraction of full-length transmembrane proteins from membranes has been described (Fleig et al., 2012; Garza et al., 2009), it is also conceivable that the eight-pass transmembrane protein EVI/WLS is cleaved within the ER membrane and that the parts are extracted separately. It remains questionable if proteins might be removed from the ER membrane by brute force generated by VCP alone and a hitherto undiscovered channel protein and/or potential cleaving enzymes seem a more elegant and potentially less energy-intensive approach.

We reported previously that EVI/WLS is a target of regulatory ERAD and that the interaction of EVI/WLS with WNT ligands prevented its degradation. For this analysis we used HEK293T cells, a cell line with low endogenous expression of WNT ligands, to demonstrate the stabilisation of EVI/WLS by overexpressing WNT ligands (Glaeser et al., 2018). Here, we show that this effect can be reversed by inhibiting the lipidation of endogenous WNT ligands in melanoma cells which naturally produce a lot of endogenous WNT5A (Figure 5B). Surprisingly, we found that inhibiting the proteasome led to the accumulation of ubiquitinated EVI/WLS even in cells with endogenous WNT ligands, indicating that cells have a surplus production of EVI/WLS leading to constant turn-over in cells with active WNT signalling (Figure 5D).

Many cancer entities require EVI/WLS and active WNT secretion throughout tumourigenesis (Zhan et al., 2017). Besides WNT signalling, the deregulation of various other cellular communication pathways also leads to cancer development and many of the underlying mechanisms are closely associated with the UPS (Deng et al., 2020). Thus, it is not surprising that FAF2 and ERLIN2 have also been implicated in cancer, for example uveal melanoma or breast cancer, respectively (Li et al., 2018; G. Wang et al., 2012). It will be interesting to test if the observed phenotypes are connected to EVI/WLS protein abundance and whether WNT signalling and tumour invasiveness could be targeted *via* ERAD.

## MATERIAL & METHODS

### Cell lines and culture method

The human melanoma cell line A375 (ATCC CRL-1619) and human embryonic kidney HEK293T (ATCC CRL-11268) cells were purchased from the American Type Culture Collection (ATCC). Cells were cultured as monolayers in Dulbecco‘s Modified Eagle‘s Medium (Gibco DMEM, 41965062, Thermo Fisher Scientific Inc.) with 4.5 g/l (high) glucose and L-glutamine supplemented with 10 % fetal bovine serum (volume fraction, Sigma Aldrich/Merck KGaA) without antibiotics at 37 °C and 5 % CO_2_ in a humidified atmosphere and regularly confirmed to be mycoplasma negative. The Clustered Regularly Interspaced Short Palindromic Repeats (CRISPR)/Cas9 *EVI/WLS* knock-out (KO) cell lines HEK293T KO2.9 and A375 sgEVI2_4 were generated by Oksana Voloshanenko or Iris Augustin, respectively, using the guide RNA sgEVI2 (TGGACGTTTCCCTGGCTTAC) and single cell clonal expansion according to previously published protocols (Glaeser et al., 2018).

### Inhibitor treatment

The Porcupine (PORCN) inhibitor LGK974 was used at 10 µM for 96 h before cell lysis with daily medium changes and PBS washes (stock solution: 50 mM in DMSO, WuXi AppTec). The proteasome inhibitor MG132 was used at 1 µM for 24 h (stock solution: 10 mM in DMSO, 474791, Sigma Aldrich/Merck KGaA). For all inhibitors, equivalent volumes of DMSO were used as solvent control.

### siRNA transfection / RNAi experiments

Cells were transfected with siRNAs from Ambion (5 µM stock solution, Thermo Fisher Scientific Inc.) or Dharmacon (20 µM stock solution, Horizon Discovery Ltd.) using Invitrogen Lipofectamine RNAiMAX Transfection Reagent (13778150, Thermo Fisher Scientific Inc.) and harvested 72 h later. All references of siRNAs are listed in Appendix Table S2.

7×10^4^ A375 melanoma cells in 2 ml culture medium per well of a 6-well plate were transfected 24 h after seeding after being washed once with phosphate buffered saline (PBS). 3 µl of siRNA or 6 µl RNAiMAX were each mixed with 125 µl of Gibco Roswell Park Memorial Institute (RPMI) 1640 Medium (11875093, Thermo Fisher Scientific Inc.) and incubated separately at room temperature for 2 min. Then, the two volumes were combined and incubated at room temperature for additional 5 min before being added to the cells. For TUBE assays, 5×10^5^ A375 cells in 10 ml culture medium were transfected in 10 cm dishes using 20 µl siRNA or 40 µl RNAiMAX in 625 µl RPMI 1640 medium each.

HEK293T cells were reverse transfected with siRNA stock solutions diluted 1/40 in ddH_2_O (working solution). Per well of a 6-well plate, 4 µl RNAiMAX were diluted in 250 µl RPMI 1640 medium and incubated at room temperature for 10 min, then further diluted with 250 µl RPMI 1640 medium. In parallel, 100 µl of siRNA working solution were added to the well and then mixed with 500 µl of the diluted RNAiMAX. 3×10^5^ HEK293T cells were seeded per well in 1.4 ml culture medium after 30 min incubation at room temperature.

### Plasmid generation and transfection

ERLIN1-FLAG (pENTR #18722573, open), ERLIN2-FLAG (pENTR #127630018, open), FAF2-FLAG (pENTR #191683255, open), UBXN4-FLAG (pENTR #178534864, open), and FLAG-UBE2K (pENTR #123919860, open) were generated using the Gateway Cloning system with the destination vectors pDEST-FLAG N-terminal (#1121, FLAG-UBE2K) and pDEST-FLAG C-terminal (#1124, all others). A STOP codon was introduced at the end of the coding sequence of FLAG-UBE2K using the Q5 Site-Directed Mutagenesis Kit (E0554S, New England Biolabs GmbH) according to the manufacturer’s instructions and the primers TGATTGGACCCAGCTTTCTTG and GTTACTCAGAAGCAATTCTG.

PORCN-FLAG was obtained from OriGene Technologies Inc. (pCMV6-Myc-DDK-tagged PORCN, #RC223764). pRK5-HA-Ubiquitin constructs (#17608, #17606, #17605, (Lim et al., 2005), #22901, (Livingston et al., 2009)), pLX302 Luciferase-V5 puro (#47553, (Kang et al., 2013)) and pcDNA Wnt3 (#35909, (Najdi et al., 2012)) were obtained from Addgene.

pcDNA-NanoLuc-Wnt3 plasmid was generated based on pcDNA Wnt3 by introducing NanoLuciferase (Hall et al., 2012) sequence after amino acid p.W26.

pcDNA-V5-hWls K410/419R (EVI/WLS-V5 K410/419R) was derived from pcDNA-V5-hWls (EVI/WLS-V5, Belenkaya et al, 2008) using the QuikChange II Site-Directed Mutagenesis Kits (200523, Agilent Technologies Inc.) according to the manufacturer’s instructions and the primers CGGAACATCAGTGGGAGGCAGTCCAGCCTGCCAGCTATGAGCAGAGTCCGGCGGC and GCCGCCGGACTCTGCTCATAGCTGGCAGGCTGGACTGCCTCCCACTGATGTTCCG. For plasmid transfection, 5×10^5^ A375 cells or 3.5×10^6^ HEK293T cells were seeded in 10 ml culture medium in 10 cm dishes the day before the transfection. The next day, 1.5 µg plasmid DNA was diluted in 500 µl serum-free RPMI 1640 and supplemented with 12 µl TransIT-LT1 Transfection Reagent (731-0027, Mirus Bio LLC). After 15 min incubation at room temperature, the mixture was added drop-wise to the cells. Cells were harvested 48 h after transfection. For 6 wells, 7×10^4^ A375 cells or 2×10^5^ HEK293T cells were seeded in 2 ml culture medium and transfected with 1 µg plasmid DNA in 250 µl RPMI 1640 medium and 5 µl transfection reagent (unless otherwise indicated).

### Reverse-transcription quantitative PCR (RT-qPCR)

For mRNA expression analysis, total RNA was isolated from cultured cells using the RNeasy Mini Kit (74106, QIAGEN GmbH) with on-column DNase digestion with the RNase-Free DNase Set (79254, QIAGEN GmbH), both according to the manufacturer’s instructions (quick start protocol version: March 2016, including optional centrifugation step at full speed). cDNA synthesis was performed in 1.5 ml tubes using the RevertAid H Minus First Strand cDNA Synthesis Kit (K1632, Thermo Fisher Scientific Inc.) with 1 µg to 5 µg of total RNA input and oligo (dT)_18_ primers following the manufacturer’s instructions, before being diluted to 5 ng/µl to 10 ng/µl with ddH_2_O. mRNA expression was quantified in technical triplicates using RT-qPCR performed in 384-well plates on a Roche LightCycler 480 Instrument II with dual hybridisation probes from The Universal ProbeLibrary (Roche). Oligonucleotide sequences used for RT-qPCR are listed in Appendix Table S3. *GAPDH* and *SDHA, G6PD*, or *ACTB* served as reference genes and relative mRNA expression levels were calculated using the Pfaffl method with siControl, siLuciferase, or DMSO treatment as calibrators. Data analysis was done using Microsoft Excel for Microsoft 365 and R (Version 3.6.1).

### Blue Sepharose assay

Secreted WNTs were enriched from cell culture supernatants using the affinity chromatography resin Blue Sepharose 6 Fast Flow (17-0948-01, GE Healthcare) and analysed by Western blotting according to previously published protocols (Glaeser et al., 2016). In brief, 2 ml cell culture medium from a well of a 6-well plate of nearly confluent cells was collected 24 h after medium change and centrifuged at room temperature for 10 min at 8 000 × g. The supernatant was then transferred to a new tube and Triton X-100 was added to a final volume fraction of 1 %. Per sample, 30 µl of Blue Sepharose 6 Fast Flow resin was washed twice in washing buffer (50 mM Tris-HCl, pH 7.5; 150 mM KCl; volume fraction of 1 % Triton X-100 in ddH_2_O) by centrifugation and decanting of supernatant (3 min, 2 800 × g). Then, resin was distributed equally to all samples and incubated overnight at 4 °C in a tube rotator. The following day, resin was washed 2 to 3 times as above, until the wash buffer was clear. After the last wash, resin was taken up in 200 µl 1× sodium dodecyl sulfate (SDS) buffer. The samples were boiled at 95 °C for 5 min and 40 µl of the sample used for SDS polyacrylamide gel electrophoresis (SDS-PAGE).

### Immunoprecipitation

To investigate protein interactions within the ERAD pathway, FLAG-tagged proteins were overexpressed in HEK393T wild type and *EVI/WLS* KO cells in 10-cm dishes and protein lysates harvested 48 h later in 600 µl eukaryotic lysis buffer (20 mM Tris-HCl, pH 7.4; 130 mM NaCl; 2 mM EDTA; glycerol at a volume fraction of 10 %, supplemented with protease inhibitor and Triton X-100 at a volume fraction of 1 %). To investigate EVI/WLS ubiquitination, pRK5-HA-Ubiquitin constructs (Ubiquitin wild type, K11, K48, K63) were overexpressed in A375 wild type and *EVI/WLS* KO cells in 10-cm dishes and protein lysates harvested 72 h later in eukaryotic lysis buffer. K11, K48, or K63 ubiquitin constructs with HA tag can only make linkages of the indicated types. After cell harvest, complete cell lysis was ensured by incubation on a tube rotator at 4 °C for 30 min, and protein lysates were clarified by centrifugation at full speed for 20 min in a table-top centrifuge at 4 °C. Protein content was quantified using the Pierce bicinchoninic acid (BCA) Protein-Assay kit (23225, Thermo Fisher Scientific Inc.) and using a Mithras LB 940 Multimode Microplate Reader (Berthold Technologies). Equal amounts of protein (1 mg to 3.5 mg) were used for pull-downs. Per sample, 40 µl ANTI-FLAG M2 Affinity Gel (A2220, Merck KGaA) or 15 µl Monoclonal Anti-HA-Agarose (A2095, Merck KGaA) and equal amounts of Control Agarose Beads (negative control, UM400, LifeSensors) were washed twice with 750 µl lysis buffer without Triton X-100 (centrifugation in between for 30 s at 5 000 × g), then blocked for 1 h with 2.5 % bovine serum albumin (BSA, mass fraction)/ Tris-buffered saline with Tween-20 (TBST; 10× TBST contains 1,37 M NaCl, 200 mM Tris-HCl, pH 7.6, and 1 % of volume fraction Tween-20) at 4 °C on a tube rotator (only for FLAG beads) and washed again twice as previously. The resin was then equally distributed to all samples and incubated overnight at 4 °C in a tube rotator. The next day, the resin was washed 4 to 7 times as previously, and proteins were eluted using 100 µl of 150 ng/µl 3× FLAG Peptide (F4799, Merck KGaA) or HA peptide (HY-P0239, Hölzel Diagnostika) and incubation for 30 min at 4 °C in a tube rotator, according to the manufacturer’s instructions. Then, samples were centrifuged as previously and 100 µl of the supernatant were transferred to a new tube. Pull-down samples or input controls (ca. 15 µg protein of the original clarified lysates) were prepared for SDS-PAGE by combining them with 1/5 volume fraction of 5× SDS buffer (312.5 mM Tris-HCl, pH 6.8; 0.5 M Dithiothreitol, DTT; mass fraction of 10 % SDS, and 0.1 % bromphenol blue; volume fraction of 10 % Tris-(2-carboxyethyl)-phosphin, TCEP, and 50 % glycerol) and 5 min incubation at 95 °C.

### Tandem Ubiquitin Binding Entity (TUBE) assays

Ubiquitinated proteins were analysed using TUBE2 (Agarose, 30 µl per sample, UM402, LifeSensors), TUBE1 (magnetic beads, 15 µl per sample, UM401M, LifeSensors), or K48 and K63 TUBE (FLAG, UM607 or UM604, LifeSensors), according to the manufacturer’s instructions with Control Agarose Beads as negative control. For buffer compositions, protein isolation protocol, and elution using 3× FLAG Peptide see section on ‘immunoprecipitation’. Pull-downs were done from one 10-cm dish per condition with 5×10^5^ A375 cells seeded 1 day before the start of drug or siRNA treatment. Cell harvest was done in eukaryotic lysis buffer with Triton X-100 at a volume fraction of 1 %, protease inhibitor, 5 mM N-ethylmaleimide, and 2 mM 1,10-phenanthroline (and 250 nM FLAG K48 or K63 TUBE if applicable) and was followed by protein quantification. Then, all samples were adjusted to contain the same amount of protein (0.5 mg to 1 mg), diluted with lysis buffer with inhibitors to 0.1 % Triton X-100 and incubated with TUBEs overnight on a tube rotator at 4 °C (TUBE1 and TUBE2) or pre-incubated with 250 nM FLAG K48 or K63 TUBE for 2 h before overnight incubation with 15 µl ANTI-FLAG M2 Affinity Gel or Control Agarose Beads at 4 °C on a tube rotator (FLAG K48 or K63 TUBE). The next day, samples were washed 4× in eukaryotic lysis buffer without Triton X-100 but with inhibitors and proteins were eluted using 3× FLAG Peptide (FLAG K48 or K63 TUBE) or by taking up beads in 100 µl 1× SDS buffer and boiled at 95 °C for 5 min (TUBE1 and TUBE2). For input controls, 40 µg of the original protein lysates were diluted to 200 µl with ddH_2_O and prepared for SDS-PAGE with 50 µl 5× SDS buffer by boiling for 5 min at 95 °C.

### SDS-PAGE and Western blotting

Total cellular protein lysates were isolated using eukaryotic lysis buffer (immunoprecipitations and TUBE assays) or 8 M urea/phosphate buffered saline (PBS, all other assays) and prepared for SDS-PAGE as described in section ‘immunoprecipitation’. 15 µg to 30 µg of protein lysate were used per sample harvested in 8 M urea/PBS. SDS-PAGE was performed using Invitrogen Bolt 4-12% Bis-Tris Plus Gels (NW04120BOX, NW04122BOX, or NW04125BOX Thermo Fisher Scientific Inc.) in 1× running buffer with 3-(N-morpholino)propanesulfonic acid (MOPS; 20× running buffer: 1 M MOPS, 1 M Tris-Base, 20 mM ethylenediaminetetraacetic acid, EDTA, 69.3 mM SDS in ddH_2_O) and the PageRuler Prestained Protein Ladder (26617, Thermo Fisher Scientific Inc.). Proteins were transferred to Amersham Protran NC Nitrocellulose-Membranes (10600002, Cytiva) by wet blotting in 1× transfer buffer (20× transfer buffer: 500 mM Bicine, 500 mM Bis-Tris, 20 mM EDTA) with 10 % methanol. Membranes were blocked in 5 % skim milk/TBST (mass fraction) for 30 min at room temperature and then incubated with primary antibodies overnight at 4 °C. All antibodies and dilutions are listed in Appendix Table S4. The next day, membranes were washed three times 7 min in TBST on a shaker and incubated with horseradish peroxidase (HRP)-coupled secondary antibodies for 1 h at room temperature (see Appendix Table S4 for references and dilutions) and again washed as before. Then, membranes were incubated with enhanced chemiluminescence (ECL) substrates and the HRP induced light signals were captured using Amersham Hyperfilm ECL (GE28-9068-36, Cytiva) and made visible using the COMPACT 2 NDT (PROTEC GmbH) developing machine. Immobilon Western HRP Substrate (WBKLS0100, Merck KGaA) was used for standard applications and SuperSignal West Femto Maximum Sensitivity Substrate (34095, Thermo Fisher Scientific Inc.) was used if stronger signal amplification was necessary.

### NanoLuciferase-WNT3 secretion assay (for reagent references see also sections on siRNA or plasmid transfection)

Luciferase assays were performed in a 384-well format using white, flat-bottom polystyrene plates (781073, Greiner Bio-One) with at least 7 technical replicates per biological replicate. On day 1, ca. 2 500 HEK293T cells in 50 µl culture medium were reverse transfected using 5 µl of 0.2 µM (Dharmacon) or 0.05 µM (Ambion) siRNA and 0.05 µl RNAiMAX in 10 µl serum-free RPMI 1640 medium per well. The next day, cells were additionally transfected using 0.1 µl TranIT-LT1 with 1 ng NanoLuciferase-WNT3 and 5 ng Firefly-luciferase for normalisation in 10 µl serum-free RPMI 1640 medium per well. 48 h later, the plate was centrifuged for 2 min at 650 × g and 20 µl of the medium were transferred to a second plate to measure NanoLuciferase-WNT3 in the supernatant. NanoLuciferase-WNT3 activity in supernatant and cell lysates and Firefly-Luciferase signals in the cell lysates were detected using the Promega Nano-Glo (N1130) or Firefly Luciferase Assay Systems and a Mithras LB 940 Multimode Microplate Reader (Berthold Technologies). Luminescence signals in the supernatant were normalised to NanoLuciferase signals and Firefly-luciferase signals in the cell lysates.

## AUTHOR CONTRIBUTIONS

L.W. designed, performed, and analysed experiments; A.L. and J.H. performed and analysed experiments; L.W. and M.B. designed the study; L.W. and M.B. wrote the manuscript, all authors contributed to editing the manuscript.

## ACKNOWLEDGEMENTS

We are grateful to K. Gläser, M. Holzem, D. Kranz, S. Redhai, O. Voloshanenko, T. Zhan, and other members of the Boutros laboratory for helpful discussions and critical comments on the manuscript. We would also like to thank T. Sommer and members of the Sommer laboratory for helpful discussions and comments on the project. We are also thankful to O. Voloshanenko for providing the pcDNA-NanoLuc-Wnt3 plasmid. We also thank the DKFZ Genomics and Proteomics Core Facility for providing plasmids and for providing access to instruments. L.W. was supported in part by the Deutsche Forschungsgemeinschaft (DFG, German Research Foundation) – Project number 259332240 / RTG 2099. Work in the laboratory of M.B. is in part supported by the SFB1324 on Mechanisms and Functions of Wnt signaling.

## CONFLICT OF INTEREST

The authors declare that they have no conflict of interest.

## Notes

### Competing Interest Statement

The authors have declared no competing interest.

