## Supplemental Information for "EVI/WLS function is regulated by ubiquitination and linked to ER-associated degradation by ERLIN2"

### SUPPLEMENTARY FIGURES

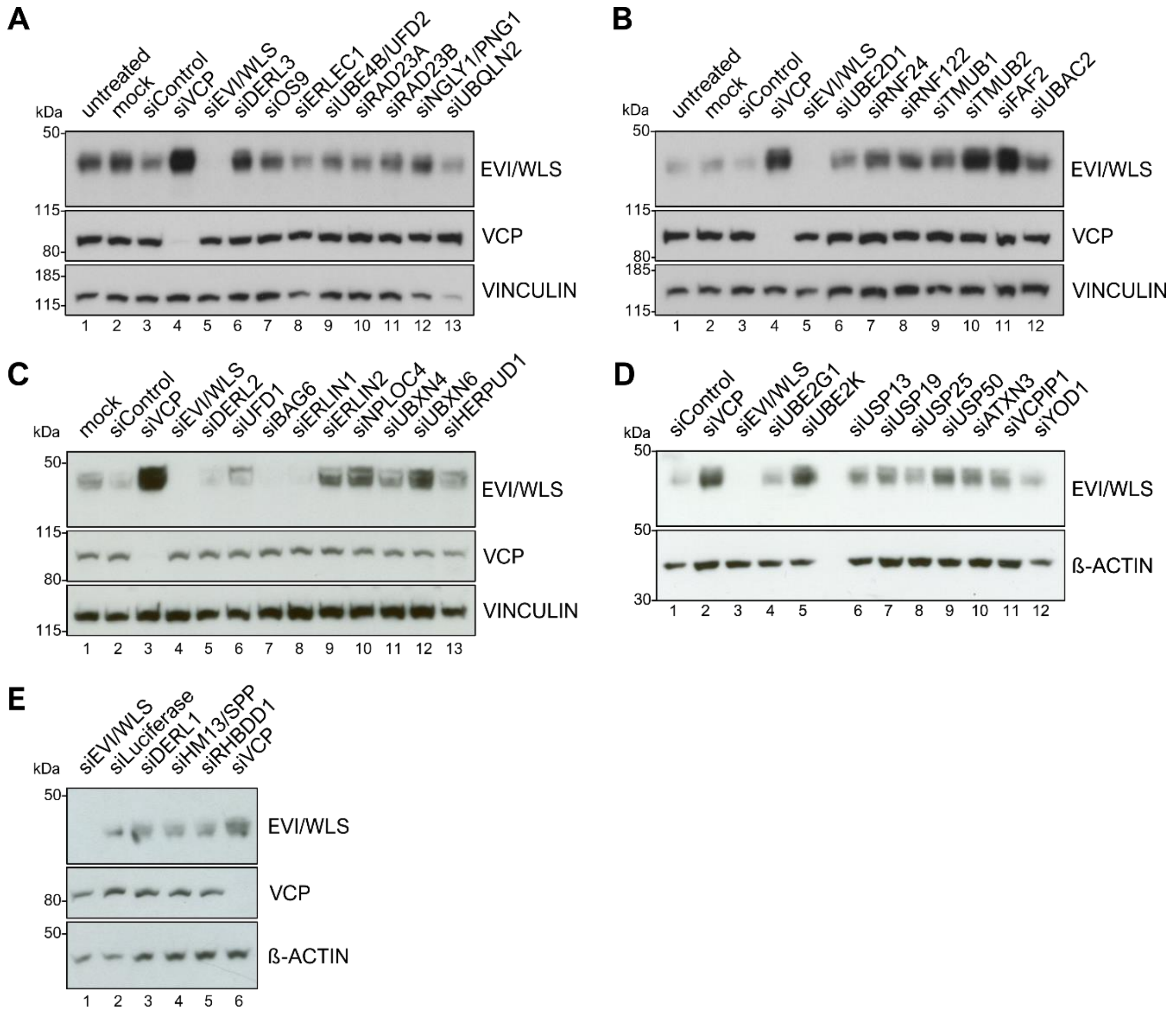

**Supplementary Figure 1.**

**siRNA-based mini-screen identifies novel candidates involved in the degradation of EVI/WLS in HEK293T cells**

EVI/WLS protein levels were analysed after siRNA mediated knock-down of target genes. Increased EVI/WLS protein levels compared to siControl/siLuciferase treatment indicated the candidate's possible involvement in EVI/WLS's ERAD process. HEK293T cells were treated with the indicated siRNAs for 72 h. VINCULIN or β-ACTIN served as loading controls. Western blots are representative of three independent experiments. kDa = kilodalton

Lanes 1-5 of **D** are also shown in main Figure 1 and were not cut to maintain integrity of the blot.

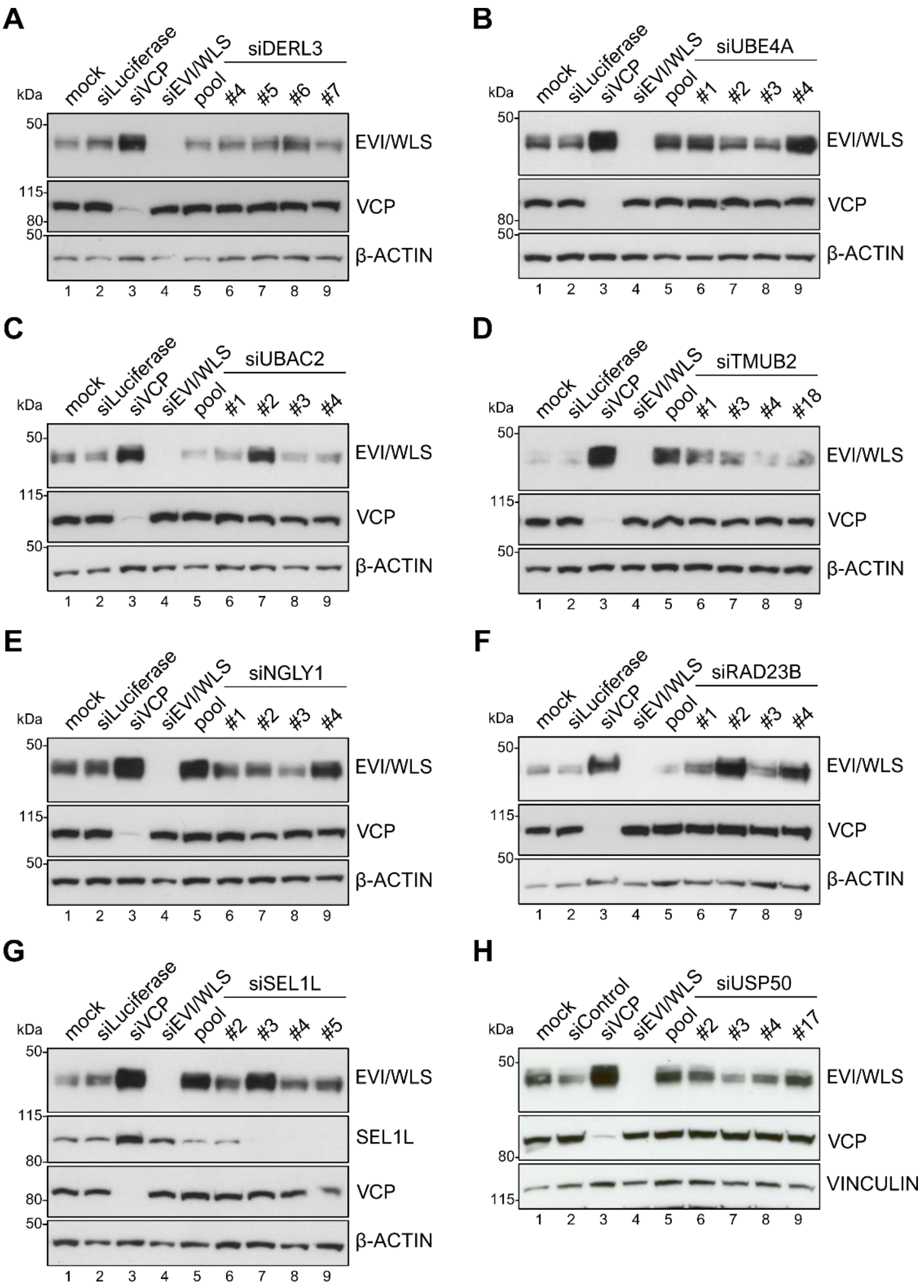

**Supplementary Figure S2. The knock-down of DERL3, UBE4A, UBAC2, TMUB2, NGLY1, RAD23B, SEL1L, or USP50 by single siRNAs did not show consistent upregulation of EVI/WLS**

EVI/WLS protein levels were not markedly elevated or varied between biological replicates after treatment with single or pooled siRNAs against the investigated candidates. HEK293T cells were treated with the indicated siRNAs for 72 h. Each gene's mRNA was targeted by either single siRNAs or an equimolecular mix of all four respective siRNAs (pool) to analyse their effect on EVI/WLS protein level. VINCULIN or β-ACTIN served as loading controls. Western blots are chosen from three independent experiments. kDa = kilodalton

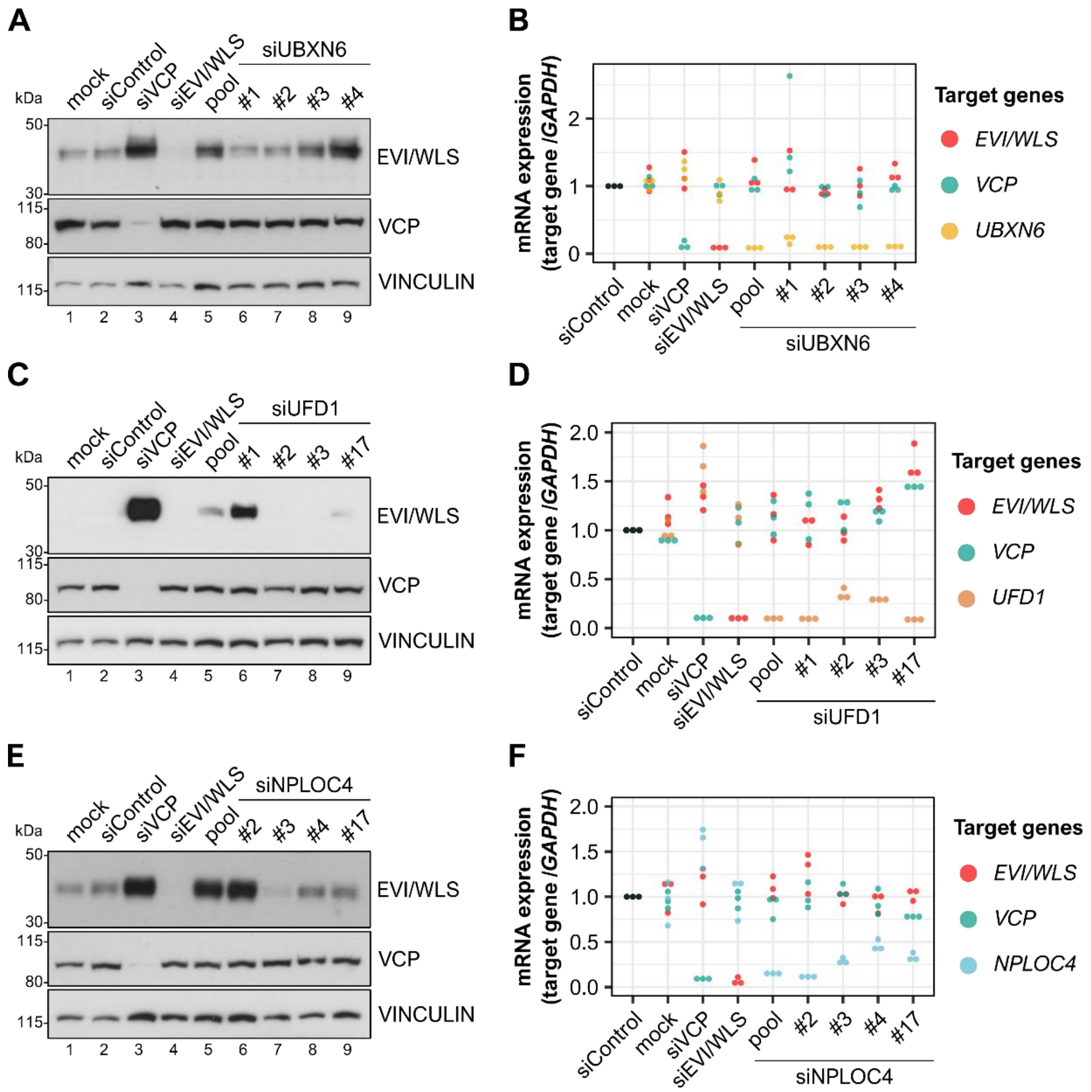

**Supplementary Figure S3.**

**The knock-down of UBXN6, UFD1, or NPLOC4 by single siRNAs did not show consistent upregulation of EVI/WLS**

EVI/WLS protein levels were not markedly elevated or varied between biological replicates after treatment with single or pooled siRNAs against the candidates investigated here (**A, C, E**). mRNA expression analyses demonstrated mostly efficient gene silencing by pooled or single siRNAs with little effects on other investigated mRNAs (**B, D, F**).

HEK293T cells were treated with the indicated siRNAs for 72 h. Each gene's mRNA was targeted by either single siRNAs or an equimolecular mix of all four respective siRNAs (pool) to analyse their effect on EVI/WLS protein level or mRNA expression. **A, C, E.** Total cell lysates were analysed by SDS-PAGE and Western blotting for the specified proteins. VINCULIN served as loading control. Western blots are representative of three independent experiments. kDa = kilodalton

**B, D, F.** Total cellular RNA was transcribed to cDNA and used for mRNA expression analyses by RT-qPCR. Target gene expression was normalised to siControl treatment and *GAPDH* served as reference gene. Individual data points from three independent experiments are shown.

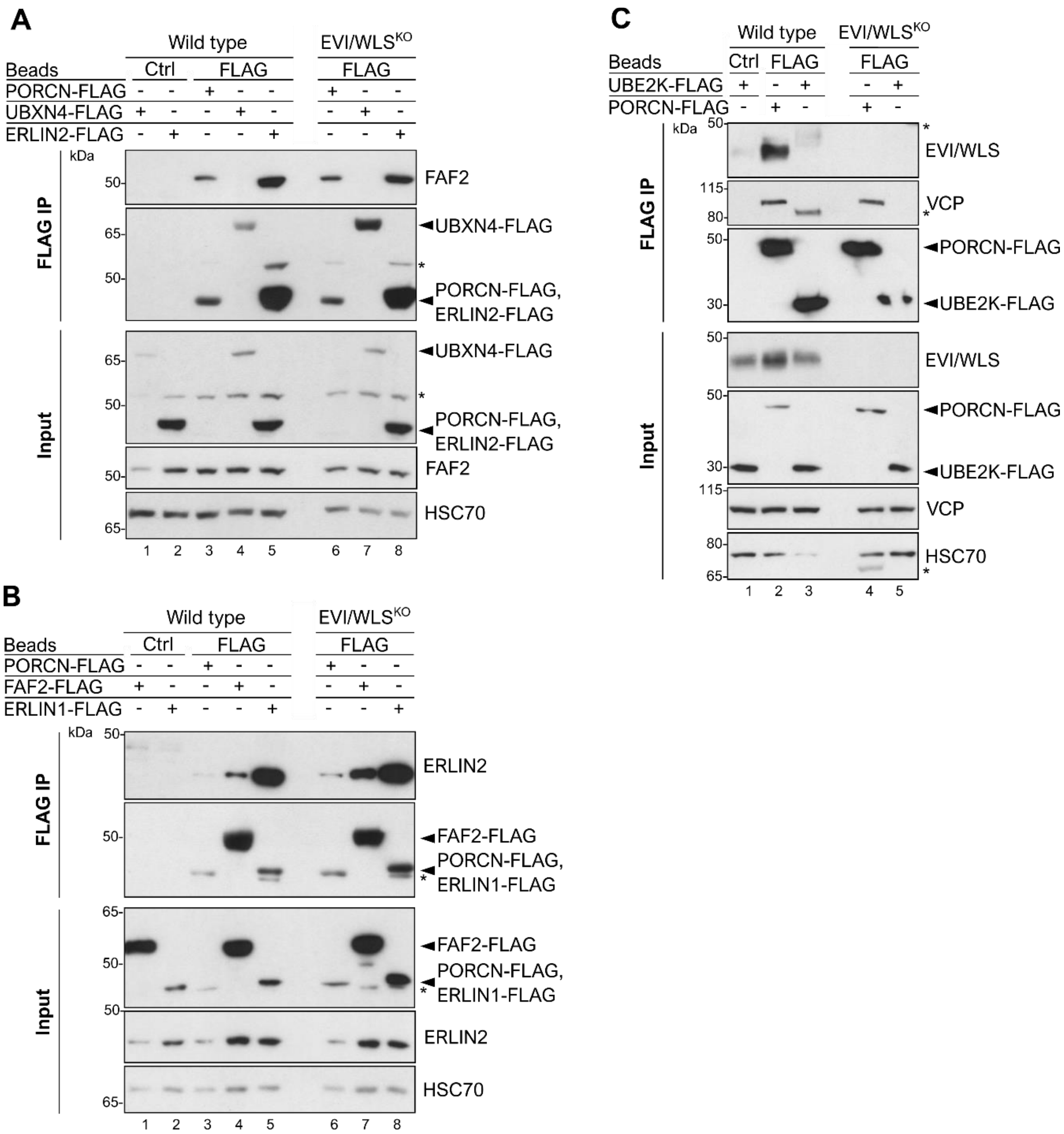

**Supplementary Figure S4. FAF2 and ERLIN2 interact with each other, but UBE2K and EVI/WLS do not**

**A, B.** IP experiments confirmed endogenous FAF2 and ERLIN2 interact with PORCN-FLAG and each other. Furthermore, endogenous ERLIN2 interacts with ERLIN1-FLAG. HEK293T wild type and EVI/WLS knock-out (EVI/WLS$^{KO}$) cells were transfected with UBXN4-FLAG, ERLIN1-FLAG, ERLIN2-FLAG, FAF2-FLAG, or PORCN-FLAG overexpression plasmids. After 48 h, total cell lysates were sampled for input control or used for FLAG IP to precipitate FLAG-tagged proteins and their interaction partners. HSC70 served as loading control. Asterisks mark signal from previous stainings. Results are representative of three independent experiments.

**C.** The binding of endogenous EVI/WLS to UBE2K-FLAG could not be detected by immunoprecipitation (IP). HEK293T wild type and EVI/WLS knock-out (EVI/WLS$^{KO}$) cells were transfected with UBE2K-N-FLAG or PORCN-FLAG overexpression plasmids. After 48 h, total cell lysates were sampled for input control or used for FLAG IP to precipitate FLAG-tagged proteins and their interaction partners. HSC70 served as loading control. Asterisks mark nonspecific signals. Results are representative of two independent experiments.

kDa = kilodalton

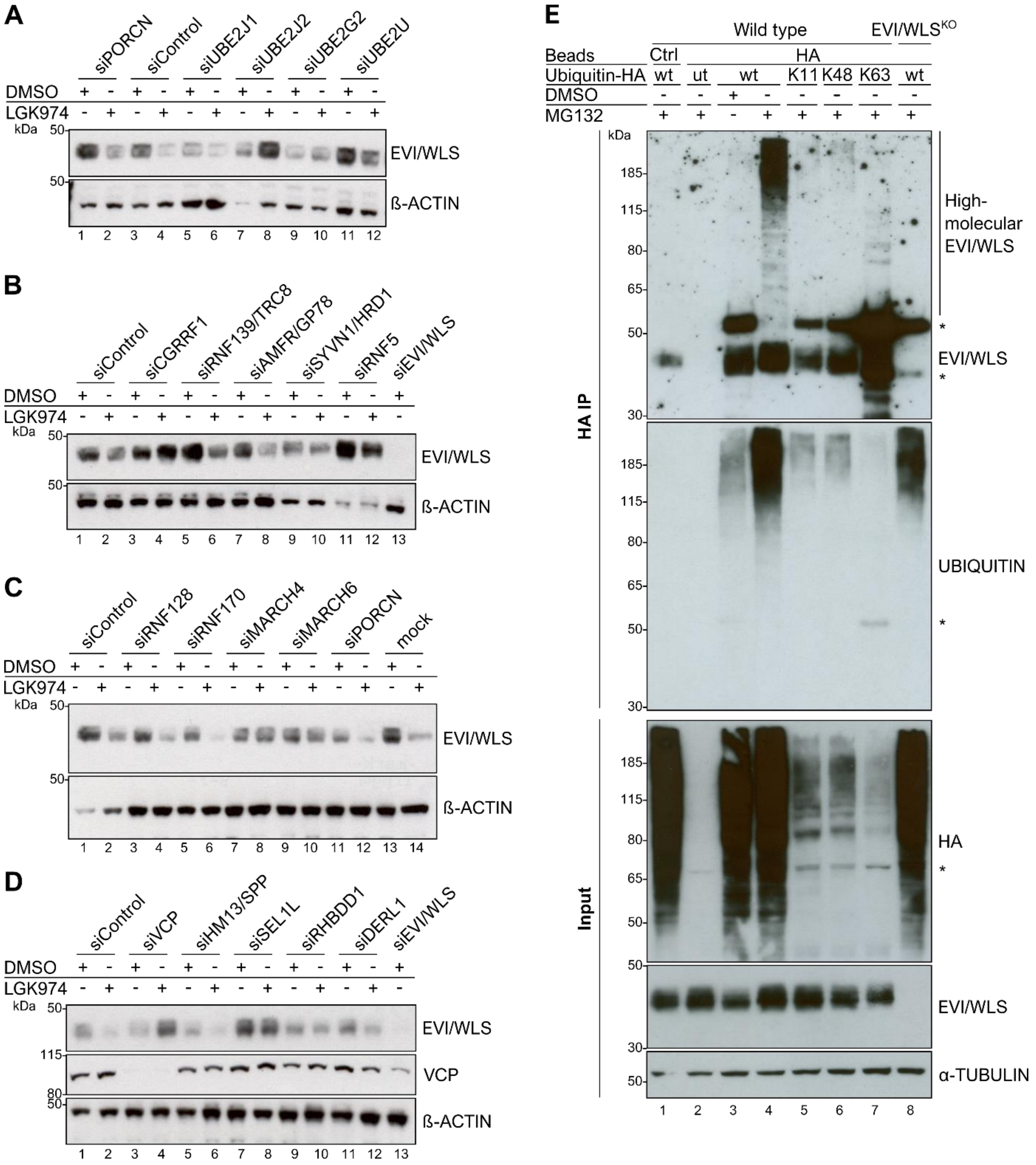

**Supplementary Figure S5. EVI/WLS is modified with K11-, K48-, and K68-linked ubiquitin and degraded with the help of VCP, CGRRF1, and UBE2J2 in A375 melanoma cells**

**A, B, C, D.** Knock-down of VCP, CGRRF1, or UBE2J2 prevented degradation of EVI/WLS after LGK974 treatment. A375 melanoma cells were treated with LGK974 (10 µM) with daily medium changes. 24 h after addition of LGK974 cells were transfected with the indicated siRNAs for 72 h. Again 72 h later, total cell lysates were analysed by SDS-PAGE and Western blotting for the specified proteins. β-ACTIN served as loading control.

**E**. Endogenous EVI/WLS is modified with ubiquitin linked via K11, K48, and K63. A375 wild type (wt) and EVI/WLS knock-out (EVI/WLS$^{KO}$) cells were transfected with pRK5-HA-Ubiquitin wt, K11, K48, or K63 overexpression plasmids or left untreated (ut). The K11, K48, and K63 ubiquitin constructs can only be elongated with further ubiquitins at the specified position, all others have been mutated to arginines. 24 h before harvest, samples were treated with the proteasome inhibitor MG132 (1 µM) or equivalent volume of DMSO as solvent control. After 72 h, total cell lysates were sampled for input control or used for HA immunoprecipitation to analyse proteins modified with HA-tagged ubiquitin. TUBULIN served as loading control. Asterisks mark nonspecific signals.

**A, B, C, D, E** Western blots are representative of three independent experiments. kDa = kilodalton

### SUPPLEMENTARY TABLES

**Table S1: Summarised results of the mini-screen**

Observed phenotypes on EVI/WLS protein abundance in HEK293T and A375 cells after knock-down of indicated target genes. Data derived from this study or from Glaeser et al., 2018; PMID: 29378775; Ref.: reference;
- no effect; + weak or variable effects; ++ strong upregulation

| | Gene | Abbreviation | NCBI gene ID | UniProt KB | tested cell line | Re-sult | Ref. |
|---|---|---|---|---|---|---|---|
| **Substrate recognition** | ERLEC1 | Endoplasmic reticulum lectin 1 | 27248 | Q96DZ1 | HEK293T | -/+ | this work |
| | ERLIN1/SPFH1 | ER Lipid Raft Associated 1 | 10613 | O75477 | HEK293T | - | this work |
| | ERLIN2/SPFH2 | ER Lipid Raft Associated 2 | 11160 | O94905 | HEK293T | +/ ++ | this work |
| | OS9/ERLEC2 | OS9 Endoplasmic Reticulum Lectin | 10956 | Q13438 | HEK293T | - | this work |
| | SEL1L/HRD3 | SEL1L Adaptor Subunit Of ERAD E3 Ubiquitin Ligase | 6400 | Q9UBV2 | HEK293T | + | this work |
| | | | | | A375 | + | this work |
| **E2 ubiquitin conjugating enzymes** | UBE2D1 | Ubiquitin Conjugating Enzyme E2 D1 | 7321 | P51668 | HEK293T | - | this work |
| | UBE2G1 | Ubiquitin Conjugating Enzyme E2 G1 | 7326 | P62253 | HEK293T | - | this work |
| | UBE2G2 | Ubiquitin Conjugating Enzyme E2 G2 | 7327 | P60604 | HEK293T | - | Glaeser et al., 2018 |
| | | | | | A375 | -/+ | this work |
| | UBE2J1 | Ubiquitin Conjugating Enzyme E2 J1 | 51465 | Q9Y385 | HEK293T | - | Glaeser et al., 2018 |
| | | | | | A375 | - | this work |
| | UBE2J2 | Ubiquitin Conjugating Enzyme E2 J2 | 118424 | Q8N2K1 | HEK293T | ++ | Glaeser et al., 2018 |
| | | | | | A375 | ++ | this work |
| | UBE2K | Ubiquitin Conjugating Enzyme E2 K | 3093 | P61086 | HEK293T | ++ | this work |
| | UBE2N/UBC13 | Ubiquitin Conjugating Enzyme E2 N | 7334 | P61088 | HEK293T | +/ ++ | this work |
| | UBE2U | Ubiquitin Conjugating Enzyme E2 U | 148581 | Q5VVX9 | HEK293T | - | Glaeser et al., 2018 |
| | | | | | A375 | - | this work |
| **E3 ubiquitin ligases** | AMFR/GP78/RNF45 | Autocrine Motility Factor Receptor | 267 | Q9UKV5 | HEK293T | -/+ | Glaeser et al., 2018 |
| | | | | | A375 | + | this work |
| | CGRRF1/RNF197 | Cell Growth Regulator With Ring Finger Domain 1 | 10668 | Q99675 | HEK293T | ++ | Glaeser et al., 2018 |
| | | | | | A375 | ++ | this work |
| | HRD1/SYVN1 | Synoviolin 1 | 84447 | Q86TM6 | HEK293T | - | Glaeser et al., 2018 |
| | | | | | A375 | - | this work |
| | MARCH4/RNF174 | Membrane Associated Ring-CH-Type Finger 4 | 57574 | Q9P2E8 | HEK293T | - | Glaeser et al., 2018 |
| | | | | | A375 | +/ ++ | this work |
| | MARCH6/TEB4/ RNF176 | Membrane Associated Ring-CH-Type Finger 6 | 10299 | O60337 | HEK293T | + | Glaeser et al., 2018 |
| | | | | | A375 | -/+ | this work |
| | RNF5 | Ring Finger Protein 5 | 6048 | Q99942 | HEK293T | -/+ | Glaeser et al., 2018 |
| | | | | | A375 | + | this work |

| | Gene | Abbreviation | NCBI gene ID | UniProt KB | tested cell line | Re-sult | Ref. |
|---|---|---|---|---|---|---|---|
| **E3 ubiquitin ligases** | RNF24 | Ring Finger Protein 24 | 11237 | Q9Y225 | HEK293T | -/+ | this work |
| | RNF122 | Ring Finger Protein 122 | 79845 | Q9H9V4 | HEK293T | -/+ | this work |
| | RNF128 | Ring Finger Protein 128 | 79589 | Q8TEB7 | HEK293T | - | Glaeser et al., 2018 |
| | | | | | A375 | -/+ | this work |
| | RNF139/TRC8 | Ring Finger Protein 139 | 11236 | Q8WU17 | HEK293T | + | Glaeser et al., 2018 |
| | | | | | A375 | - | this work |
| | RNF170 | Ring Finger Protein 170 | 81790 | Q96K19 | HEK293T | - | Glaeser et al., 2018 |
| | | | | | A375 | - | this work |
| | UBE4B/UFD2 | Ubiquitination Factor E4B | 10277 | O95155 | HEK293T | - | this work |
| **Retrotrans-location/Dis-location** | DERL1 | Derlin 1 | 79139 | Q9BUN8 | HEK293T | + | this work |
| | | | | | A375 | - | this work |
| | DERL2 | Derlin 2 | 51009 | Q9GZP9 | HEK293T | + | this work |
| | DERL3 | Derlin 3 | 91319 | Q96Q80 | HEK293T | +/ ++ | this work |
| | FAF2/ETEA/UBXD8 | Fas Associated Factor Family Member 2 | 23197 | Q96CS3 | HEK293T | +/ ++ | this work |
| | HM13/SPP | Histocompatibility Minor 13/ Signal Peptide Peptidase | 81502 | Q8TCT9 | HEK293T | + | this work |
| | | | | | A375 | - | this work |
| | NPLOC4/NPL4 | NPL4 Homolog, Ubiquitin Recognition Factor | 55666 | Q8TAT6 | HEK293T | +/ ++ | this work |
| | RHBDD1/RHBDL4 | Rhomboid Domain Containing 1 | 84236 | Q8TEB9 | HEK293T | + | this work |
| | | | | | A375 | - | this work |
| | UBAC2 | UBA Domain Containing 2 | 337867 | Q8NBM4 | HEK293T | + | this work |
| | UBXN4/ERASIN/ UBXD2 | UBX Domain Protein 4 | 23190 | Q92575 | HEK293T | +/ ++ | this work |
| | UBXN6/UBXD1 | UBX Domain Protein 6 | 80700 | Q9BZV1 | HEK293T | +/ ++ | this work |
| | UFD1/UFD1L | Ubiquitin Recognition Factor In ER Associated Degradation 1 | 7353 | Q92890 | HEK293T | +/ ++ | this work |
| | VCP/P97/CDC48 | Valosin Containing Protein | 7415 | P55072 | HEK293T | ++ | Glaeser et al., 2018 |
| | | | | | A375 | ++ | this work |

| | Gene | Abbreviation | NCBI gene ID | UniProt KB | tested cell line | Re-sult | Ref. |
|---|---|---|---|---|---|---|---|
| **Delivery to the proteasome** | BAG6/BAT3/SCYTHE | BAG Cochaperone 6 | 7917 | P46379 | HEK293T | - | this work |
| | HERPUD1/HERP | Homocysteine Inducible ER Protein With Ubiquitin Like Domain 1 | 9709 | Q15011 | HEK293T | + | this work |
| | NGLY/PNG1 | N-Glycanase 1 | 55768 | Q96IV0 | HEK293T | + | this work |
| | RAD23A | RAD23 Homolog A, Nucleotide Excision Repair Protein | 5886 | P54725 | HEK293T | -/+ | this work |
| | RAD23B | RAD23 Homolog B, Nucleotide Excision Repair Protein | 5887 | P54727 | HEK293T | -/+ | this work |
| | TMUB1/HOPS | Transmembrane And Ubiquitin Like Domain Containing 1 | 83590 | Q9BVT8 | HEK293T | - | this work |
| | TMUB2 | Transmembrane And Ubiquitin Like Domain Containing 2 | 79089 | Q71RG4 | HEK293T | +/ ++ | this work |
| | UBQLN2/DSK2 | Ubiquilin 2 | 29978 | Q9UHD9 | HEK293T | - | this work |
| **De-ubiquitin-ating enzymes** | ATXN3 | Ataxin 3 | 4287 | P54252 | HEK293T | - | this work |
| | USP13 | Ubiquitin Specific Peptidase 13 | 8975 | Q92995 | HEK293T | - | this work |
| | USP19 | Ubiquitin Specific Peptidase 19 | 10869 | O94966 | HEK293T | + | this work |
| | USP25 | Ubiquitin Specific Peptidase 25 | 29761 | Q9UHP3 | HEK293T | - | this work |
| | USP50 | Ubiquitin Specific Peptidase 50 | 373509 | Q70EL3 | HEK293T | +/ ++ | this work |
| | VCPIP1 | Valosin Containing Protein Interacting Protein 1 | 80124 | Q96JH7 | HEK293T | - | this work |
| | YOD1/OTUD2 | YOD1 Deubiquitinase | 55432 | Q5VVQ6 | HEK293T | - | this work |

**Table S2: siRNA references used in this study**

| Target gene | siRNA number | Manufacturer | Reference |
|---|---|---|---|
| EVI/WLS | 1 | Ambion | s36745 |
| EVI/WLS | 3 | Ambion | s36747 |
| siGENOME Non-Targeting siRNA Pool #1 | / | Dharmacon | D-001206-13-20 |
| siLuciferase/ RLuc Duplex siRNA | / | Dharmacon | P-002070-01-20 |
| AMFR/GP78 | 1 | Dharmacon | D-006522-01 |
| | 2 | Dharmacon | D-006522-02 |
| | 3 | Dharmacon | D-006522-03 |
| | 4 | Dharmacon | D-006522-04 |
| ATXN3 | 1 | Dharmacon | D-012013-01 |
| | 2 | Dharmacon | D-012013-02 |
| | 3 | Dharmacon | D-012013-03 |
| | 4 | Dharmacon | D-012013-04 |
| BAG6 | 1 | Dharmacon | D-005062-01 |
| | 2 | Dharmacon | D-005062-02 |
| | 3 | Dharmacon | D-005062-03 |
| | 4 | Dharmacon | D-005062-04 |
| CGRRF1 | 1 | Dharmacon | D-006933-01 |
| | 2 | Dharmacon | D-006933-02 |
| | 3 | Dharmacon | D-006933-03 |
| | 4 | Dharmacon | D-006933-04 |
| DERL1 | 2 | Dharmacon | D-010733-02 |
| | 3 | Dharmacon | D-010733-03 |
| | 4 | Dharmacon | D-010733-04 |
| | 18 | Dharmacon | D-010733-18 |
| DERL2 | 1 | Dharmacon | D-010576-01 |
| | 2 | Dharmacon | D-010576-02 |
| | 3 | Dharmacon | D-010576-03 |
| | 4 | Dharmacon | D-010576-04 |
| DERL3 | 4 | Dharmacon | D-032237-04 |
| | 5 | Dharmacon | D-032237-05 |
| | 6 | Dharmacon | D-032237-06 |
| | 7 | Dharmacon | D-032237-07 |
| ERLIN1 | 2 | Dharmacon | D-015639-02 |
| | 18 | Dharmacon | D-015639-18 |
| | 19 | Dharmacon | D-015639-19 |
| | 20 | Dharmacon | D-015639-20 |
| ERLIN2/SPFH2 | 2 | Dharmacon | D-017943-02 |
| | 3 | Dharmacon | D-017943-03 |
| | 4 | Dharmacon | D-017943-04 |
| | 5 | Dharmacon | D-017943-05 |
| FAF2/ETEA/UBXD8 | 1 | Dharmacon | D-010649-01 |
| | 3 | Dharmacon | D-010649-03 |
| | 4 | Dharmacon | D-010649-04 |
| | 17 | Dharmacon | D-010649-17 |
| HERPUD1/HERP | 1 | Dharmacon | D-020918-01 |
| | 2 | Dharmacon | D-020918-02 |
| | 3 | Dharmacon | D-020918-03 |
| | 4 | Dharmacon | D-020918-04 |
| HM13/SPP | 4 | Dharmacon | D-005896-4 |
| | 5 | Dharmacon | D-005896-5 |
| | 6 | Dharmacon | D-005896-6 |
| | 19 | Dharmacon | D-005896-19 |
| MARCH4 | 2 | Dharmacon | D-023172-02 |
| | 3 | Dharmacon | D-023172-03 |
| | 5 | Dharmacon | D-023172-05 |
| | 6 | Dharmacon | D-023172-06 |

| Target gene | siRNA number | Manufacturer | Reference |
|---|---|---|---|
| MARCH6 | 1 | Dharmacon | D-006925-01 |
| | 2 | Dharmacon | D-006925-02 |
| | 3 | Dharmacon | D-006925-03 |
| | 4 | Dharmacon | D-006925-04 |
| NGLY/PNG1 | 1 | Dharmacon | D-016457-01 |
| | 2 | Dharmacon | D-016457-02 |
| | 3 | Dharmacon | D-016457-03 |
| | 4 | Dharmacon | D-016457-04 |
| NPLOC4/NPL4 | 2 | Dharmacon | D-020796-02 |
| | 3 | Dharmacon | D-020796-03 |
| | 4 | Dharmacon | D-020796-04 |
| | 17 | Dharmacon | D-020796-17 |
| PORCN | 1 | Dharmacon | D-009613-01 |
| | 2 | Dharmacon | D-009613-02 |
| | 3 | Dharmacon | D-009613-03 |
| | 4 | Dharmacon | D-009613-04 |
| RAD23B | 1 | Dharmacon | D-011759-01 |
| | 2 | Dharmacon | D-011759-02 |
| | 3 | Dharmacon | D-011759-03 |
| | 4 | Dharmacon | D-011759-04 |
| RHBDD1/RHBDL4 | 1 | Dharmacon | D-019378-01 |
| | 2 | Dharmacon | D-019378-02 |
| | 3 | Dharmacon | D-019378-03 |
| | 4 | Dharmacon | D-019378-04 |
| RNF128 | 1 | Dharmacon | D-007061-01 |
| | 4 | Dharmacon | D-007061-04 |
| | 17 | Dharmacon | D-007061-17 |
| | 18 | Dharmacon | D-007061-18 |
| RNF139/TRC8 | 1 | Dharmacon | D-006942-01 |
| | 2 | Dharmacon | D-006942-02 |
| | 4 | Dharmacon | D-006942-04 |
| | 17 | Dharmacon | D-006942-17 |
| RNF170 | 1 | Dharmacon | D-007078-01 |
| | 2 | Dharmacon | D-007078-02 |
| | 3 | Dharmacon | D-007078-03 |
| | 4 | Dharmacon | D-007078-04 |
| RNF5 | 1 | Dharmacon | D-006558-01 |
| | 2 | Dharmacon | D-006558-02 |
| | 3 | Dharmacon | D-006558-03 |
| | 18 | Dharmacon | D-006558-18 |
| SEL1L | 2 | Dharmacon | D-004885-02 |
| | 3 | Dharmacon | D-004885-03 |
| | 4 | Dharmacon | D-004885-04 |
| | 5 | Dharmacon | D-004885-05 |
| SYFN/HRD1 | 1 | Dharmacon | D-007090-01 |
| | 2 | Dharmacon | D-007090-02 |
| | 3 | Dharmacon | D-007090-03 |
| | 4 | Dharmacon | D-007090-04 |
| TMUB2 | 1 | Dharmacon | D-014307-01 |
| | 3 | Dharmacon | D-014307-03 |
| | 4 | Dharmacon | D-014307-04 |
| | 18 | Dharmacon | D-014307-18 |
| UBAC2 | 1 | Dharmacon | D-107914-01 |
| | 2 | Dharmacon | D-107914-02 |
| | 3 | Dharmacon | D-107914-03 |
| | 4 | Dharmacon | D-107914-04 |
| UBE2G1 | 1 | Dharmacon | D-010154-01 |
| | 2 | Dharmacon | D-010154-02 |
| | 4 | Dharmacon | D-010154-04 |
| | 5 | Dharmacon | D-010154-05 |

| Target gene | siRNA number | Manufacturer | Reference |
|---|---|---|---|
| UBE2G2 | 1 | Dharmacon | D-009095-01 |
| | 2 | Dharmacon | D-009095-02 |
| | 3 | Dharmacon | D-009095-03 |
| | 5 | Dharmacon | D-009095-05 |
| UBE2J1 | 1 | Dharmacon | D-007266-01 |
| | 3 | Dharmacon | D-007266-03 |
| | 19 | Dharmacon | D-007266-19 |
| | 20 | Dharmacon | D-007266-20 |
| UBE2J2 | 1 | Dharmacon | D-008614-01 |
| | 2 | Dharmacon | D-008614-02 |
| | 4 | Dharmacon | D-008614-04 |
| | 18 | Dharmacon | D-008614-18 |
| UBE2K | 17 | Dharmacon | D-009431-17 |
| | 18 | Dharmacon | D-009431-18 |
| | 19 | Dharmacon | D-009431-19 |
| | 20 | Dharmacon | D-009431-20 |
| UBE2N | 1 | Dharmacon | D-003920-01 |
| | 2 | Dharmacon | D-003920-02 |
| | 4 | Dharmacon | D-003920-04 |
| | 5 | Dharmacon | D-003920-05 |
| UBE2U | 1 | Dharmacon | D-008998-01 |
| | 2 | Dharmacon | D-008998-02 |
| | 3 | Dharmacon | D-008998-03 |
| | 4 | Dharmacon | D-008998-04 |
| UBE2V1 | 2 | Dharmacon | D-010064-02 |
| | 21 | Dharmacon | D-010064-21 |
| | 22 | Dharmacon | D-010064-22 |
| | 23 | Dharmacon | D-010064-23 |
| UBE2V2 | 1 | Dharmacon | D-008823-01 |
| | 2 | Dharmacon | D-008823-02 |
| | 3 | Dharmacon | D-008823-03 |
| | 4 | Dharmacon | D-008823-04 |
| UBE4A | 1 | Dharmacon | D-007200-01 |
| | 2 | Dharmacon | D-007200-02 |
| | 3 | Dharmacon | D-007200-03 |
| | 4 | Dharmacon | D-007200-04 |
| UBXN4/ERASIN/UBXD2 | 3 | Dharmacon | D-014184-03 |
| | 4 | Dharmacon | D-014184-04 |
| | 17 | Dharmacon | D-014184-17 |
| | 18 | Dharmacon | D-014184-18 |
| UBXN6/UBXD1 | 1 | Dharmacon | D-008785-01 |
| | 2 | Dharmacon | D-008785-02 |
| | 3 | Dharmacon | D-008785-03 |
| | 4 | Dharmacon | D-008785-04 |
| UFD1L/UFD1 | 2 | Dharmacon | D-017918-02 |
| | 3 | Dharmacon | D-017918-03 |
| | 4 | Dharmacon | D-017918-04 |
| | 17 | Dharmacon | D-017918-17 |
| USP13 | 1 | Dharmacon | D-006064-01 |
| | 2 | Dharmacon | D-006064-02 |
| | 3 | Dharmacon | D-006064-03 |
| | 4 | Dharmacon | D-006064-04 |
| USP19 | 3 | Dharmacon | D-006068-03 |
| | 4 | Dharmacon | D-006068-04 |
| | 5 | Dharmacon | D-006068-05 |
| | 6 | Dharmacon | D-006068-06 |
| USP25 | 2 | Dharmacon | D-006074-02 |
| | 3 | Dharmacon | D-006074-03 |
| | 4 | Dharmacon | D-006074-04 |
| | 5 | Dharmacon | D-006074-05 |

| Target gene | siRNA number | Manufacturer | Reference |
|---|---|---|---|
| USP50 | 2 | Dharmacon | D-031837-2 |
| | 3 | Dharmacon | D-031837-3 |
| | 4 | Dharmacon | D-031837-4 |
| | 17 | Dharmacon | D-031837-17 |
| VCP | 5 | Dharmacon | D-008727-05 |
| | 6 | Dharmacon | D-008727-06 |
| | 7 | Dharmacon | D-008727-07 |
| | 8 | Dharmacon | D-008727-08 |
| VCPIP1 | 1 | Dharmacon | D-019137-01 |
| | 2 | Dharmacon | D-019137-02 |
| | 3 | Dharmacon | D-019137-03 |
| | 4 | Dharmacon | D-019137-04 |
| YOD1 | 1 | Dharmacon | D-027369-01 |
| | 2 | Dharmacon | D-027369-02 |
| | 3 | Dharmacon | D-027369-03 |
| | 4 | Dharmacon | D-027369-04 |

**siRNAs from Dharmacon Genomewide 96 well plates MTP**

| Target gene | siRNA number | Manufacturer | Reference |
|---|---|---|---|
| ERLEC1 | / | Dharmacon | M-010658-00 |
| OS9 | / | Dharmacon | M-010811-00 |
| RAD23A | / | Dharmacon | M-005231-00 |
| RNF122 | / | Dharmacon | M-007068-00 |
| RNF24 | / | Dharmacon | M-006943-00 |
| TMUB1 | / | Dharmacon | M-018578-00 |
| TMUB2 | / | Dharmacon | M-014307-00 |
| UBAC2 | / | Dharmacon | M-017914-00 |
| UBE2D1 | / | Dharmacon | M-009387-01 |
| UBQLN2 | / | Dharmacon | M-013566-00 |
| UFD2/UBE4B | / | Dharmacon | M-007202-01 |

**Table S3: Sequences of primers used for RT-qPCR**

| Target mRNA | Forward primer sequence (5' -> 3') | Reverse primer sequence (5' -> 3') | Probe | Probe Reference |
|---|---|---|---|---|
| ACTB | CCAACCGCGAGAAGATGA | CCAGAGGCGTACAGGGATAG | #64 | 4688635001 |
| AXIN2 | GCTGACGGATGATTCCATGT | ACTGCCCACACGATAAGGAG | #56 | 4688538001 |
| ERLIN2/SPFH2 | GGAAGAAGGCGCTCATTG | TGAAATCTTCTCAGTCTCCTTC | #29 | 4687612001 |
| EVI/WLS | TCATGGTATTTCAGGTGTTTCG | GCATGAGGAACTTGAACCTAAA A | #38 | 4687965001 |
| FAF2/ETEA/ UBXD8 | GAAGGAGGAGGAGGTGCAA | TCCTTTCCTTTTCCTCCTGTAA | #82 | 4689054001 |
| GAPDH | AGCCACATCGCTCAGACAC | GCCCAATACGACCAAATCC | #60 | 4688589001 |
| NPLOC4 | CGGTTTACATCAATAGAAACAAGAC T | AACAACAAATCGCCATGCTT | #25 | 4686993001 |
| PORCN | GCTACTGCAAGGCTGTCTCC | GCTTCAGGTAGGATGGCAAC | #3 | 4685008001 |
| SDHA | GGACCTGGTTGGTCTTTGGTC | CCAGCGTTTGGTTTAATTGG | #80 | 4689038001 |
| UBE2K | AGGACCTCCAGACACACCAT | CGGACCTTAGGGGGATTAAA | #69 | 4688686001 |
| UBE2N | CGCAGGATCATCAAGGAAA | AAATAACGGGCGTTGCTCT | #72 | 4688953001 |
| UBXN4/ERASIN/ UBXD2 | CGCTTCGGTGGTACTGTTG | TCCCAACAAGGTCCAAATGT | #4 | 4685016001 |
| UBXN6 | CCTGGACAACATCCACCTG | AGGCAGTTAATGCGCTCCT | #63 | 4688627001 |
| UFD1l/UFD1 | CAGCATGAGGAGTCGACAGA | CCAGTCTATTGCCAGATCCAG | #67 | 4688660001 |
| VCP | AGAGGCAGACAAACCCATCA | AGTGATCTCGACGGATCTCAG | #35 | 4687680001 |

**Table S4: Antibodies, TUBEs, and reagents for IP used in this study**

**Primary antibodies**

| Target protein | Reference | Host | Clonality | Dilution | Supplier |
|---|---|---|---|---|---|
| β-ACTIN | ab6276 | mouse | monoclonal (AC-15) | 1/ 10 000 | Abcam |
| β-ACTIN | #4967 | rabbit | polyclonal | 1/ 1 000 | Cell Signaling Technology |
| ERLIN2/SPFH2 | EB06896 | goat | polyclonal | 1/ 1 000 | VWR |
| EVI/WLS | 655902 | mouse | monoclonal (YJ5) | 1/ 250 – 1/ 1 000 | BioLegend |
| FAF2/ETEA/UBXD8 | GTX14759 | goat | polyclonal | 1/ 1 000 | GeneTex |
| FLAG | F1804 | mouse | monoclonal (M2) | 1/ 1 000 | Sigma-Aldrich |
| FLAG | F7425 | rabbit | polyclonal | 1/ 8 000 | Sigma-Aldrich |
| HA-Tag | #2367 | mouse | monoclonal (6E2) | 1/ 1 000 | Cell Signaling Technology |
| HSC70 | sc-7298 | mouse | monoclonal (clone B-6) | 1/ 2 000 | Santa Cruz Biotechnology |
| K48-linkage Specific Polyubiquitin | #8081 | rabbit | monoclonal (D9D5) | 1/ 100 000 | Cell Signaling Technology |
| K63-linkage Specific Polyubiquitin | #5621 | rabbit | monoclonal (D7A11) | 1/ 1 000 | Cell Signaling Technology |
| SEL1L | ab78298 | rabbit | polyclonal | 1/ 1 000 | Abcam |
| α-TUBULIN | #2144 | rabbit | polyclonal | 1/ 3 000 | Cell Signaling Technology |
| UBE2K/E2-25K | MAB6609 | mouse | monoclonal (701316) | 1/ 1 000 | R&D Systems |
| UBE2N | #6999 | rabbit | monoclonal (D2A1) | 1/ 1 000 | Cell Signaling Technology |
| Ubiquitin | #3936 | mouse | monoclonal (P4D1) | 1/ 3 000 | Cell Signaling Technology |
| VCP/P97 | ab11433 | mouse | monoclonal (5) | 1/ 100 000 | Abcam |
| VINCULIN | AB6039 | rabbit | polyclonal | 1/ 150 000 | Merck Millipore |
| WNT3 | GTX128100 | rabbit | polyclonal | 1/ 1 000 | GeneTex |
| WNT5A/B | #2530 | rabbit | monoclonal (C27E8) | 1/ 1 000 | Cell Signaling Technology |

**Horse-radish peroxidase (HRP)-coupled antibodies for Western blotting**

| Specificity | Reference | Host | Clonality | Conjugate | Dilution | Supplier |
|---|---|---|---|---|---|---|
| β-ACTIN | SC47778 HRP | mouse | monoclonal (C4) | HRP | 1/ 5 000 | Santa Cruz Biotechnology |
| Anti-Goat IgG | 6160-05 | rabbit | polyclonal | HRP | 1/ 5 000 | SouthernBiotech |
| Anti-Mouse IgG (H+L) | AB_10015289 | goat | polyclonal | HRP | 1/ 10 000 | Jackson ImmunoResearch |
| Anti-Rabbit IgG (H+L) | AB_2313567 | goat | polyclonal | HRP | 1/ 10 000 | Jackson ImmunoResearch |

**Tandem Ubiquitin Binding Entities (TUBEs), immunoprecipitation, and control reagents**

| Name | Specificity | Conjugate | Reference | Supplier |
|---|---|---|---|---|
| TUBE Control | Control | Agarose beads | UM400 | LifeSensors |
| TUBE1 | Pan-ubiquitin | Magnetic beads | UM401M | LifeSensors |
| TUBE2 | Pan-ubiquitin | Agarose beads | UM402 | LifeSensors |
| K48-TUBE | K48-linked ubiquitin | FLAG-tag | UM607 | LifeSensors |
| K63-TUBE | K63-linked ubiquitin | FLAG-tag | UM604 | LifeSensors |
| M2 AFFINITY GEL | Anti-FLAG | Agarose beads | A2220 | Sigma-Aldrich |
| Monoclonal Anti-HA-Agarose | Anti-HA | Agarose beads | A2095 | Sigma-Aldrich |
| BLUE SEPHAROSE 6 Fast Flow | General protein purification | Sepharose | 17-0948-01 | GE Healthcare |
